# Human iPSC-derived salivary gland organoids model diabetic salivary gland dysfunction

**DOI:** 10.1101/2025.10.01.679897

**Authors:** Devon Duron Ehnes, Akira Morishita, Ashish Phal, Hee Yun Jung, Yen Chian Lim, Zachary Foreman, Vincenzo Cirulli, Julie Mathieu, Hannele Ruohola-Baker

## Abstract

Salivary glands are highly susceptible to injury and degeneration. To facilitate studies in human salivary gland disease, we developed a rapid protocol for 3D hiPSC-derived salivary gland organoids that recapitulate human fetal salivary gland gene expression and function, have both ductal and acinar cell types, secretory capacity, and the ability to respond to cholinergic agonism. Oral health issues resulting from diabetes mellitus have been attributed to salivary gland dysfunction, leading to chronic xerostomia and increased dental caries. To study diabetic salivary gland hypofunction, we further developed a diabetic model, demonstrating diabetic hallmarks including FOXO1 nuclear localization, AGE-RAGE signaling, and defective oxidative phosphorylation, which were by treatment with the diabetic drug Metformin. Our model has implications for the development of effective therapeutics against salivary gland dysfunction in diabetes and other metabolic diseases in exocrine tissues.

## Introduction

Salivary glands are exocrine glands in the mouth that produce and secrete saliva into the oromaxillary facial region. Saliva is a critical, multifunctional fluid that not only maintains lubrication in the oral cavity, but also facilitates mastication and digestion^1,2^, swallowing^3^, and speech^2,4^, performs antimicrobial functions^5,6^, and has even been found to be a critical player in sensory perception in the mouth and mouthfeel experience of food^2,7,8^. Salivary gland dysfunction can be caused by numerous injuries, including those caused by irradiation during the treatment of cancer, and various diseases, and infections^9–12^. Despite abundant potential causes of salivary gland dysfunction, the majority of these can be attributed to immune misregulation^13–15^, age-related degeneration^16–18^, and diabetes^19–21^. According to the NIDDK, an estimated 1 in 9 Americans suffers from diabetes mellitus and is projected to increase to 1 in 6 by 2030^22^. In the scheme of diabetic studies, the oral consequences, which include oral burning, severe tooth decay, disrupted wound healing, and increased susceptibility to oral infections^23^, have been largely overlooked^24^. Research has shown that diabetes-driven dysfunction happens through various tissue-by-tissue mechanisms^25^. In diabetic human blood plasma, elevated glucose contributes to increased oxidative stress and circulating inflammatory cytokines, which leads to increased inflammatory response, oxidative stress, damaged barrier function, and cell death^26^. However, it is unknown what cell populations within a salivary gland are affected, whether they are affected by defective cell organization, function, or metabolic response.

Organoids are three-dimensional multicellular tissue-like systems derived from stem cells. Given the right cues, they can develop and self-organize to recapitulate original tissues’ architecture, functionality, and genetic signature of original tissues in vivo. Accordingly, they have become popular for modeling development and disease, screening pharmaceuticals, and studying development and morphogenesis on a large scale in vitro^27–29^ and in numerous tissue systems^30–32^ including salivary glands, though the ability to create a scalable, high-throughput model has remained elusive. The predominant existing salivary gland organoid protocols create something known as a salisphere, a 3D cluster of cells derived from primary adult salivary gland cells^33–37^. While salispheres have been useful in regenerative and developmental studies, they lack scalability necessary for high-throughput screening applications and are limited in disease modeling capacity. Recent efforts have developed organoids derived from pluripotent cells^38,39^, however, these protocols are highly technically complex, making them suboptimal for large-scale disease modeling/therapeutic screening efforts. Using organoids to study metabolic diseases is gaining particular interest due to the human-specific characteristics that cannot be recapitulated in animals^40^, namely differences in metabolism that underlie development, function, and pharmaceutical processing^41–43^. To date, organoid models have been created to recapitulate diabetic islet cells^44,45^ and vascular tissue^46^, as well as some other secondary tissues, including kidney^47^ and liver^48^, but salivary gland organoid models of diabetes and other metabolic diseases have yet to be developed.

Using our single-cell atlas of human fetal salivary gland development^49,50^, we created a rapid single-cell combinatorial indexing (sci) RNA-Seq-guided protocol to develop salivary gland organoids from human induced pluripotent stem cells. These organoids recapitulate salivary gland morphology and show transcriptional and functional similarities to native salivary glands, including the development of acinar and ductal compartments. We further developed organoids into a diabetic model that reproduces key diabetic characteristics and is responsive to pharmaceutical treatment.

## Results

### Development of salivary gland organoids from human iPSCs

We developed a protocol to differentiate human iPSCs into functional salivary glands guided by human fetal developmental cues (Figure 1A). We created a single-cell RNA-Seq atlas from which we identified critical pathways driving the development of different populations in the human fetal salivary gland^49^, the information from which we used to guide our decisions during protocol development. We explored several different parameters to optimize this protocol, described in Figures S1-S3 and supplemental text. To create 3D organoids, we began by seeding iPSCs in ultra-low attachment six-well plates. To reduce cell stress and death in the transition from 2D to 3D, we plated iPSCs at 400,000 cells per well in mTeSR media supplemented with 10µM ROCKi. This allowed cells to attach and form spheroids. Once spheroids were formed, we transitioned into the oral epithelium portion of the differentiation for a period of 10 days using a previously developed protocol^50^. We then switched to driving salivary gland fate. In generating our human single-cell salivary gland atlas, we detected cell-cell signaling interactions between support tissues and various clusters as the salivary epithelium developed. We evaluated how they changed as the tissue developed. Several FGF factors were identified (*FGF10*, *FGF1*, *FGF7*, Figure S2D-F) which are critical for controlling budding and branching in salivary gland development^51–53^. We also observed significant expression of EGF family factor *NRG1* in the mesenchyme (Figure S2G), which has only recently been identified as a driver in salivary acinar specification^54^. We observed that, except for *FGF10*, whose expression level remained relatively constant, the expression level of these factors varied by developmental stage. Therefore, using *FGF10* expression as the baseline, we determined the ratio of expression of each of these factors (Figure S2H). Using these ratios, we varied the amount of these factors that we added at different stages of organoid development to best mimic in vivo expression patterns. We capitalized on what we learned with EGF treatment in the 2D optimization experiments (Document S2), adding this factor late in differentiation to drive acinar differentiation. Previous studies have also demonstrated that treating salivary precursors with ROCKi increases proacinar identity in salispheres^37^, so we also added a short period of ROCKi treatment in the latter stages of differentiation.

**Figure 1:**
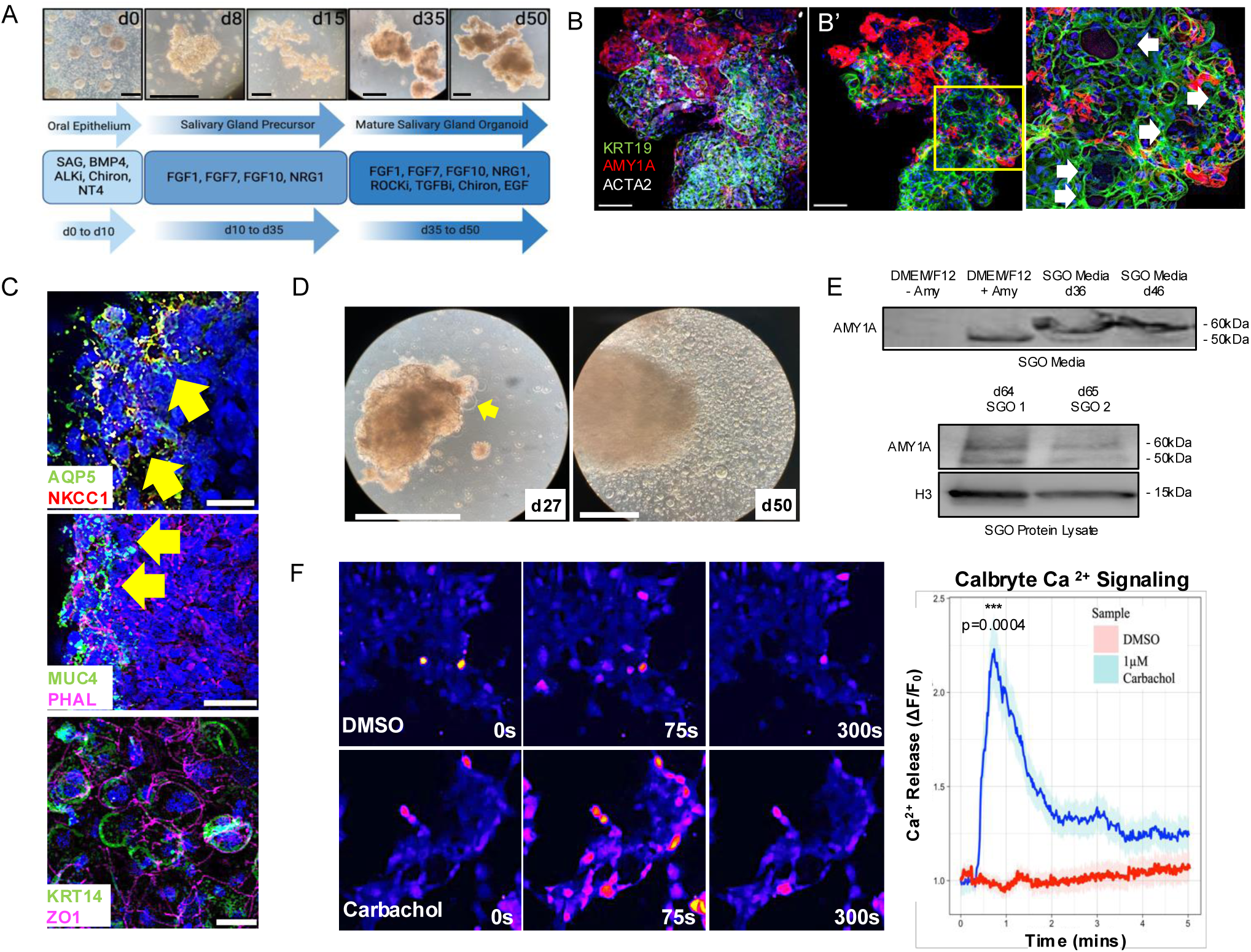
Salivary gland organoids derived from hiPSCs exhibit structural and functional similarities to salivary glands. (A) Schematic of protocol to create iPSC-derived salivary gland organoids. Scale bars = 500µm (B) d50 salivary gland organoids stained with AMY1A which marks acinar, KRT19 which marks ducts, and ACTA2 which marks myoepithelial cells. Scale bars = 50µm (B’) Subset of slices (10 slices) of B to show lumens in that region of the organoid (C) Immunofluorescence of other salivary gland markers. Scale Bars = 25µm, yellow arrows identify specified marker localized to lumens in the organoid (D) d27 and d50 salivary gland organoids producing fluid droplets. Scale bar = 500µm (E) Western blot of salivary gland organoid media shows amylase expression. (F) Evaluation of calcium response to Carbachol. Brighter pink and yellow indicates high calcium response. Salivary gland organoids exhibit significantly higher activity than DMSO controls. Scale bars = 25µm

We observed that organoids expressed salivary amylase (AMY1A) by d35 and that organoids adopt an increased degree of cellular organization by d50, wherein AMY1A+ acinar regions group separately from KRT19+ ductal regions, and are flanked by ACTA2+ myoepithelial progenitors, recapitulating in vivo salivary gland organization (Figure 1B). Organoids exhibit some heterogeneity in their shape, size, and how much AMY1A expressed. Still, we always observe the organization of distinct AMY1A+ acinar regions and KRT19+ ductal regions (Figure S3G-I”). We also found that cells within the organoids organized into lumenized epithelial structures reminiscent of the ductal networks observed in salivary glands (Figure 1B’) and that they expressed salivary markers including the lubricating glycoprotein MUC4, the sodium/potassium transporter NKCC1, and water channel protein AQP5 (Figure 1C) on the apical side of those lumen-like regions, in line with their localization and expression in vivo^55,56^. We also observe the presence of some KRT14+ progenitor cells nestled among asymmetrically localized ZO-1+ cells (Figure 1C), indicating the presence of less mature regions within the organoids^57^. After about 20 days, we observed the presence of acellular droplets (Figure 1D, yellow arrow) which exhibit a rolling, fluid-like motion when the plate is moved (Video S1), and eventually detach from the organoid and fall to the bottom of the well. These become more numerous as organoids mature. To evaluate the contents of these droplets, we collected this media at each feed between days 30-50, concentrated it, and assessed it with Coomassie staining (Figure S4B) and Western blot (Figure 1E, Figure S4C). Coomassie staining demonstrated consistent loading between samples and showed that the media contained an abundance of proteins located at the same molecular mass as salivary amylase. Probing the media for salivary amylase produced multiple bands at sizes consistent with what has previously been described in saliva^58,59^. We also assayed protein lysates of organoids from two experiments (Figure 1E, Figure S4A). We found that both experimental runs yielded amylase expression after organoids reach maturation, suggesting organoids are producing and secreting amylase in these droplets. Salivary secretion is initiated by parasympathetic signaling though cholinergic receptor activation by acetylcholine^60–63^, leading to Ca^2+^ signaling response. To determine whether our organoids could react to a similar response, we loaded cells with a fluorescent calcium indicator (Calbryte) and then either with 1µM DMSO or the cholinergic agonist carbachol. We observed that the carbachol activated calcium signaling in salivary gland organoid cells almost immediately upon treatment, exhibiting sustained activity for nearly 2 minutes, compared to DMSO controls (Figure 1F).

### iPSC-derived organoids exhibit transcriptomic similarities to human fetal salivary glands

To evaluate how efficient our differentiation is and to determine what cell populations are present in our organoids, we conducted single-cell RNA sequencing (sciRNA-Seq) on the salivary gland organoids (Figure 2A). Unbiased clustering resulted in 5 unique clusters (Figure 2B). Pseudotime analysis (Figure 2C) suggests that cells transition from the least mature cells in cluster 4 to the most mature cells in cluster 1. Transcriptomic analysis of salivary gland- and epithelium-associated markers showed that markers shifted from broader epithelial markers (KRT18, CDH2, KITLG) in the Early Epithelium cluster to more cell-type specific mature markers in the Acinar-like cluster, including the salivary gland enriched cholinergic receptor CHRM3, which leads to the activity observed in Figure 1F. Importantly, we observed that several factors that are secreted in saliva^64,65^ were highly expressed in the Acinar-like cluster (Figure 2D, Figure S5A), supporting their ability to produce a saliva-like substance. Because of the developmental and functional similarities between pancreas and salivary gland acinar cells^66^, we also assessed clusters for expression of pancreatic markers (Figure S5B) which were minimal, compared to known salivary markers (Figure S5B, blue box). To evaluate how our human fetal development-guided salivary gland organoids compared to developing human fetal tissue, we compared salivary gland organoids sci-RNA-Seq to human fetal salivary gland sci-RNA-Seq by mapping the transcriptome of salivary gland organoid cells to the original human fetal clusters (Figure 2E).

**Figure 2:**
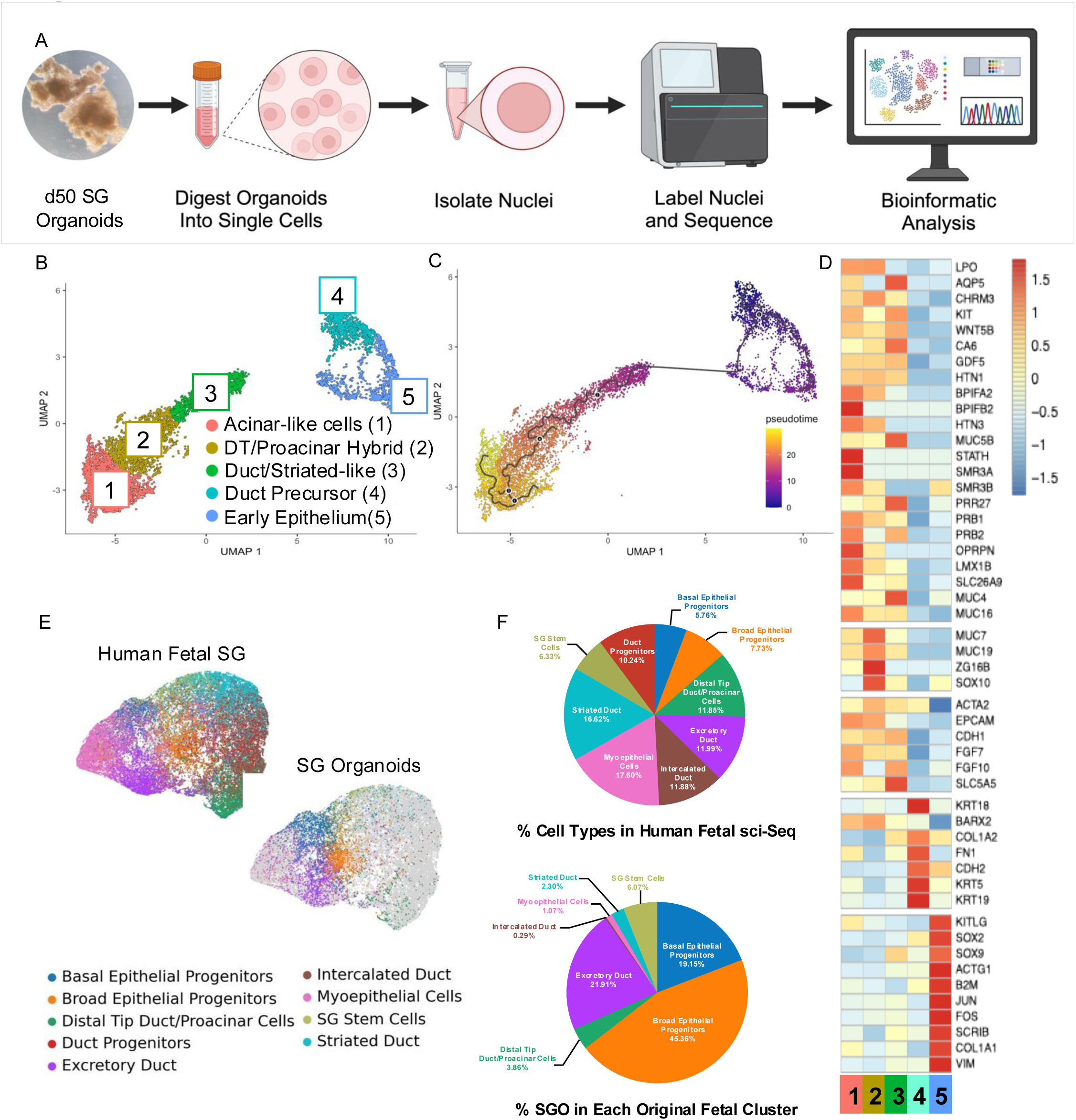
iPSC-derived salivary gland organoids show transcriptional overlap with developing human fetal salivary glands. (A) Schematic of organoid sci-RNA Seq. (B) Unbiased UMAP of organoids gives 5 clusters. (C) Pseudotime analysis shows that Cluster 5 is the earliest cluster, giving rise to the later, more mature clusters. (D) Gene expression of salivary gland markers shows earlier markers of less mature and progenitor cells are highly expressed in earlier clusters. In contrast, more mature markers and saliva components are described in later clusters. (E) Comparison of salivary gland organoids to human fetal salivary gland development shows that cells from the organoids map to all clusters in the fetal dataset. (F) Percent of cell types in each cluster or original human fetal dataset and percent of salivary gland organoid cells that map to those clusters.

In the original fetal sci-RNA-Seq analysis, we observed a relatively even distribution of salivary epithelium cell types (Figure 2F). The largest populations were myoepithelial cells (17.6%) and striated duct (16.62%), with other terminal populations comprising roughly 10-12% each, alongside smaller proportions of the progenitor groups. In our salivary gland organoid data set, the majority of the cells fall into the various progenitor cell types, a common phenomenon in organoid cultures derived from pluripotent cells^67,68^, with broad and basal epithelial progenitor types accounting for nearly 65% of the cells. However, importantly, we were able to identify all expected terminal cell types, accounting for roughly 35% of the total population, including a small population (6.07%) of salivary gland stem cells. These findings suggest that our method effectively captures all cell type bifurcations in the developmental trajectory and recapitulates the cell heterogeneity found in the salivary gland.

### Human iPSC-derived salivary gland organoids can model diabetic salivary gland dysfunction

We adapted a previously developed protocol for vascular organoids^46^ to model diabetic conditions in salivary glands. When vascular organoids were exposed to diabetic conditions (4.5-fold glucose and 1ng/ml each of TNF*α* and IL-6), organoids exhibited common diabetic vascular phenotypes. We hypothesized that using similar conditions on salivary organoids would recapitulate some responses to diabetic conditions in vitro, so we treated mature organoids for an additional two weeks in either osmotic control media or diabetic media (Figure 3A). Compared to osmotic control, diabetic salivary gland organoids exhibited reduced AMY1A and disrupted KRT19 expression (Figure 3B). Previous studies have suggested that malfunction in various tissues in diabetes mellitus result from FOXO1 nuclear localization^69,70^ which leads to defective mitochondrial function^70,71^. We assessed FOXO1 activity in osmotic control and diabetic salivary gland organoids, revealing that diabetic conditions resulted in nearly 7 times the amount of nuclear FOXO1 (Figure 3C), suggesting that defects in the salivary gland may be related to mitochondrial stress. Ultrastructural evaluation of organoids by scanning electron microscopy (Figure 3D) showed that organoids cultured under control and diabetic culture conditions both exhibit ultrastructural features and organelles reminiscent of an exocrine cell phenotype including the presence of large vacuoles filled with electron-dense proteinaceous material (Figure 3D, Black *), and secretory vesicles (Figure 3D, White *) that appear more abundant in the control samples, a phenotype that has been previously reported in murine diabetic models^72^. The most notable changes observed in organoids cultured under diabetic conditions included an increased frequency of adherens junctions with altered ultrastructural organization (Figure 3D, red arrows) that lack compaction, exhibit a clear gap (i.e., white space) between intercellular plaques at the cell-cell junction, and have increased interaction with electron-dense cytoskeletal elements. We also observed an increased frequency of autophagic bodies (Figure 3D, black arrows) and ER stress (Figure 3D, blue arrows). To evaluate the secretory capacity of the organoids in diabetic conditions, we assessed the protein and amylase in the media. We observed an increase in overall proteins in the diabetic samples, visualized with Coomassie (Figure S4B). However, western blots against AMY1A in the organoid media showed decreased AMY1A, indicating that the diabetic cells secreting less AMY1A into their media. To assess transcriptomic changes observed in diabetic conditions, we conducted bulk RNA sequencing on diabetic organoids from two independent experiments (Figure S6A, B). Compared to osmotic controls, diabetic samples exhibited comparatively few genes significantly upregulated in diabetic samples; most statistically significant genes were downregulated (Figure 3F). KEGG analysis of the upregulated genes (Figure 3G) returned terms for AGE-RAGE signaling pathway^73–75^ and protein absorption and digestion^76,77^, both of which are recognized as indicators and drivers of diabetic pathology. AGE-RAGE signaling is driven by advanced glycated end products (AGEs). These products interact with their receptor RAGE and are known to modulate several downstream stress response factors. The AGE-RAGE response is exacerbated in the diabetic environment. It is believed to create a form of metabolic memory that persists even when the diabetic environment is regulated, leading to diabetic complications^78^. Though this signaling pathway can be expressed in healthy cells in response to routine cell stress, we observe that the expression of these factors is very low in our controls and only elevated in the diabetic samples (Figure 3H). Consistent with this, we observed an increase in the expression of pathogenic COLIV in the diabetic samples (Figure 3I) that was not evident in the controls. KEGG analysis also returned Focal Adhesion as a significant term from upregulated genes. Studies have shown that hyperglycemia stimulates focal adhesion remodeling to facilitate actin dynamics and vesicle trafficking^79,80^ but that sustained activation of focal adhesion kinases induces the activation of inflammatory response signaling, exacerbating tissue damage/malfunction^81–83^. KEGG analysis of downregulated genes (Figure 3J) returned Oxidative Phosphorylation, Ribosome and Proteasome, all of which have been observed in several diabetic tissues^25,84–88^. Additionally, though hyperglycemia can initially upregulated ribosome biogenesis^89^, chronic hyperglycemia and inflammation ultimately lead to disrupted biogenesis due to nucleotide depletion^90^. Finally, the proteasome functions to remove misfolded or oxidized proteins, and several studies^91,92^ have shown decreased proteasomal activity in diabetic conditions, leading to insulin resistance, ER stress, tissue dysfunction^91,93,94^. These results indicate that our salivary gland organoid model responds to the diabetic environment in a manner consistent with that observed in other tissues. However, it does not define what causes tissue dysfunction in the salivary gland. When we evaluate how the diabetic environment impacts the expression of salivary gland markers, we observe that marker expression is largely unaffected (Figure 3K). There is some slight reduction in MUC1, a critical component in lubricating mucus, thought to act in an anti-inflammatory capacity^95^. There is also slight reduction in cystatins, which are secreted proteolytic proteins found in saliva^96,97^, and a reduction in several keratins which provide structural support in ducts and having some functional signaling^98^. Though misregulation of the expression of these genes may ultimately contribute to dysfunction, the mild change in expression indicates that their expression may not be the primary driver.

**Figure 3:**
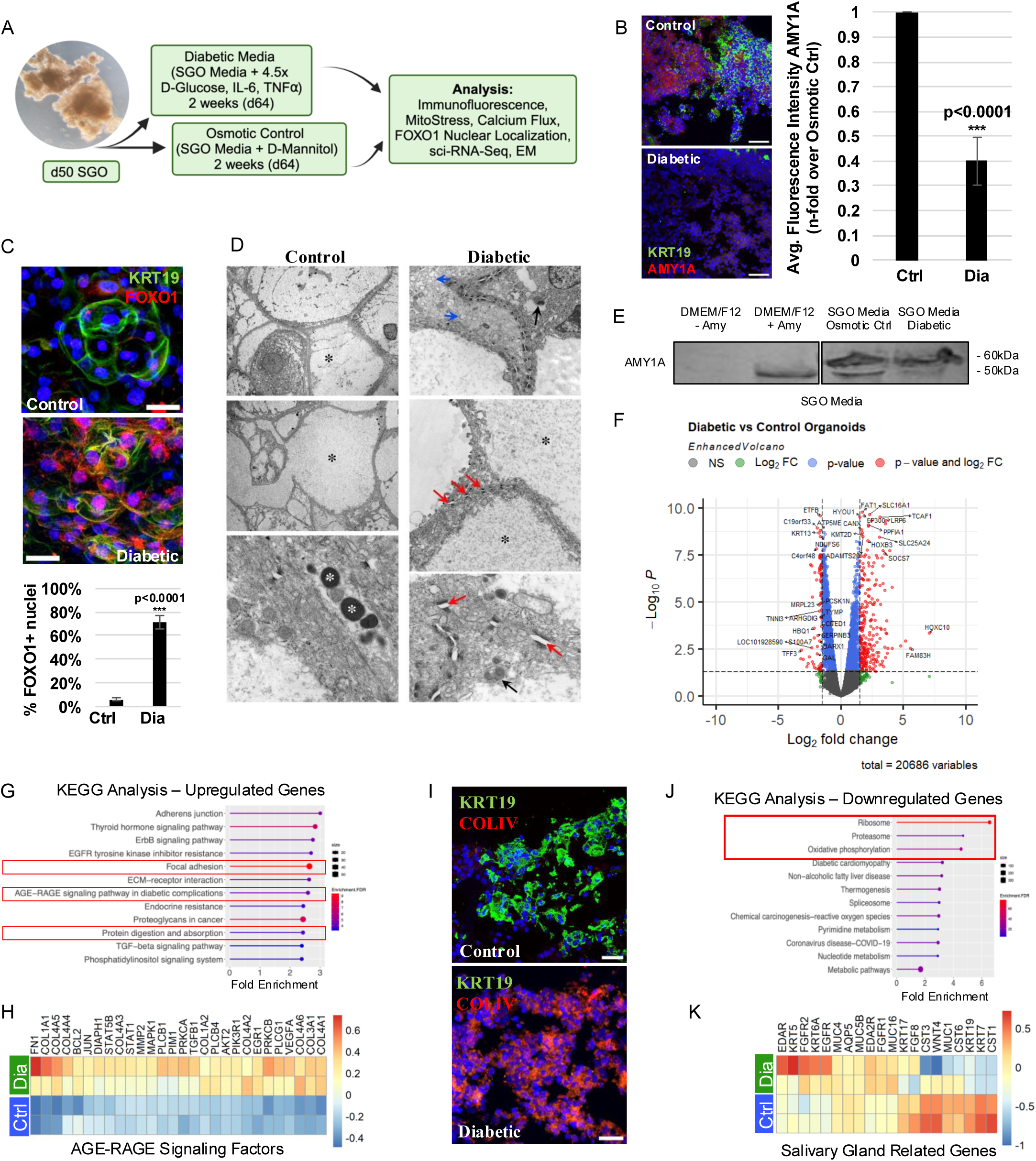
Salivary gland organoids grown in diabetic conditions exhibit signatures of diabetic tissues. (A) Schematic of diabetic organoid development. (B) Immunofluorescence of osmotic control and diabetic salivary gland organoids for AMY1A and KRT19, showing reduced AMY1A expression in diabetic samples. Results represent the average n-fold change over control for 4 individual experiments each with n=3-9. Scale bars = 50µm (C) Diabetic organoids have significantly more FOXO1+ nuclei than osmotic controls. n=5, scale bars = 25µm (D) Scanning Electron Microscopy of osmotic control and diabetic salivary gland organoids show differences in gap junctions, total protein, and autophagic bodies. (E) Western blot of organoid culture media to evaluate amylase secretion between osmotic control and diabetic organoids. (F) Volcano plot of genes up- and downregulated in bulk RNA sequencing of osmotic control and diabetic organoids. (G) KEGG analysis of genes upregulated in diabetic conditions. (H) Upregulated genes associated with AGE-RAGE signaling. (I) Expression of pathogenic COLIV in osmotic control and diabetic organoids. Scale bars = 50µm (J) KEGG analysis of genes downregulated in diabetic conditions. (K) Expression of salivary gland-related genes between osmotic control and diabetic conditions.

### hiPSC salivary gland organoid disease model shows mitochondrial dysfunction in diabetic models

Oxidative phosphorylation is a critical metabolic process that occurs in mitochondria. Further evaluation of the genes associated with disrupted oxidative phosphorylation included reduced expression of genes regulating several complexes in the electron transport chain (Sup. Figure6C, Figure4A). Still, the most prominent of these were complex I and ATPases. Complex I dysfunction is consistent with studies in other diabetic tissues that showed that diabetes-induced redox imbalance resulted in mitochondrial dysfunction through the disruption of this complex, leading to further defects in the synthesis of macromolecules including DNA, lipids, and proteins^99^. Expression changes in ATPases showed that V-type ATPases, which are critical to the function of intracellular organelles including secretory vesicles, endosomes, and lysosomes^100,101^, and which have been previously associated with exocrine secretory function^100,102^, were markedly upregulated, while F-type ATPases (ATP Synthases), were downregulated (Figure 4A, ATPases). We also evaluated the effect of the diabetic environment on gene expression of other processes regulating mitochondrial function and health. For instance, the accumulation of ROS has been recognized as a pathological driver in a variety of tissues^103–107^. However, we found that our diabetic conditions showed marginal changes in expression of a limited number of antioxidants compared to osmotic controls, including the expression of *GSS*, which synthesizes the primary antioxidant glutathione, as well as the expression of its transporter *SLC25A39* (Figure S6D). Additionally, mitochondrial fission and fusion are dynamic processes that occur in a balanced manner and help mitochondria deal with environmental stresses^108^; fusion results in the combination of two mitochondria, creating elongated mitochondrial networks, while fission results in fragmentation^109^. Interestingly, while changes in the expression of mitofusins have been observed in many other tissues, we saw no change in the expression of the mitofusins. Instead, we observed a decrease in the expression of the translocase of the outer membrane (TOMM) proteins that assists in mitochondrial fusion (Figure S6E), as well as a decrease in the expression of several inner membrane transport proteins (Figure S6F). When we evaluated the morphology of the mitochondria in salivary gland organoids, we observed that the osmotic controls exhibited many interconnected mitochondrial networks, while the diabetic organoids show fragmentation (Figure 4B-C). Using the Mitochondria Analyzer ImageJ plugin^110^, we found that the mitochondria in the diabetic samples exhibited a more spherical morphology and fewer branches per mitochondria (Figure4C), characteristics of more fragmented mitochondria that lack the extensive networks of healthy mitochondria, indicating that our diabetic conditions perturb mitochondrial fusion in the face of stress.

**Figure 4:**
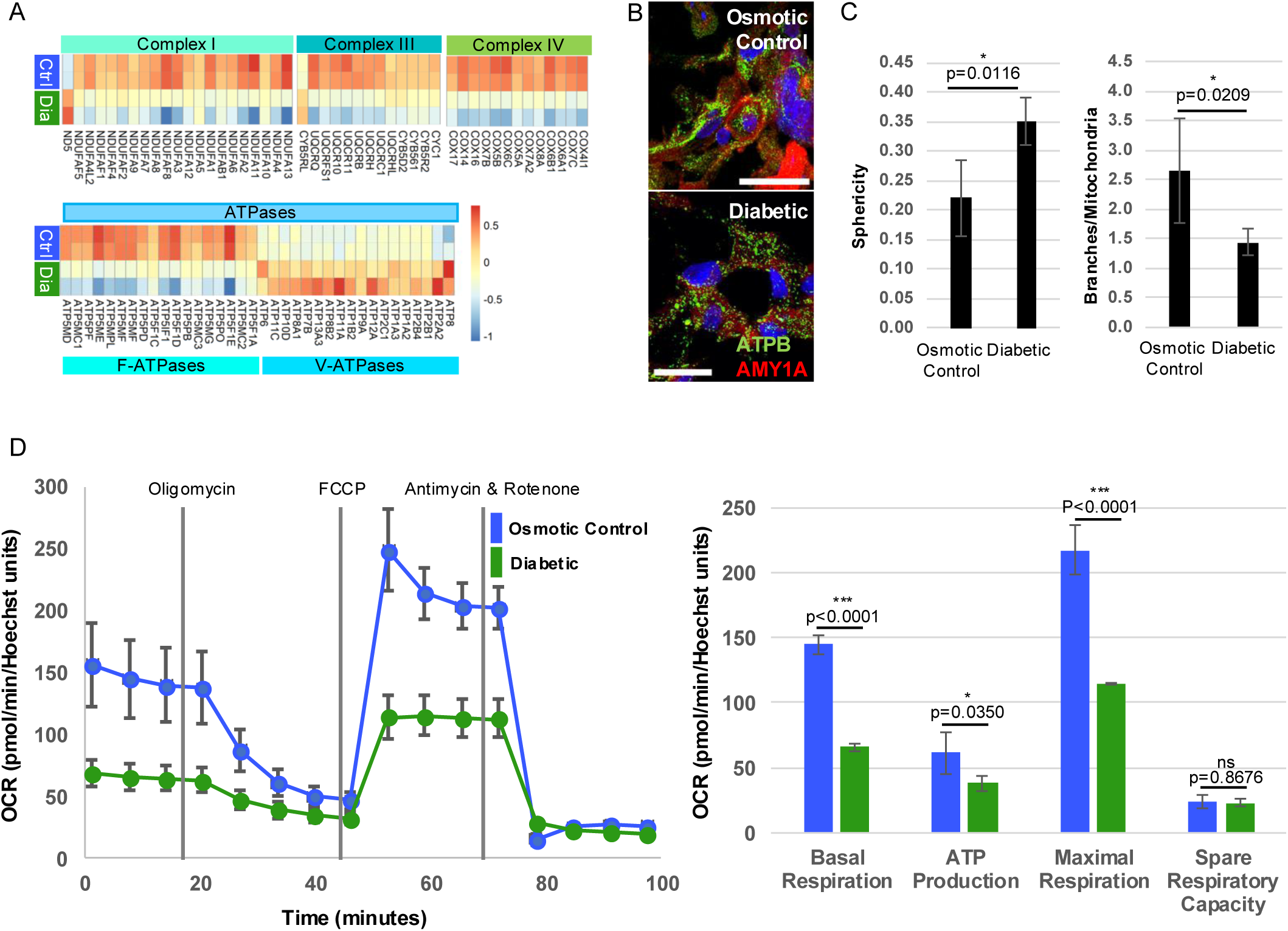
Mitochondrial activity is perturbed in diabetic salivary gland organoids. (A) Expression of classes of genes in different mitochondrial complexes between osmotic control and diabetic organoids. (B) Mitochondria morphology is different in control and diabetic organoids. n=3, scale bars = 25µm (C) Quantification of sphericity and branches per mitochondria supports increased fragmentation in diabetic organoids. (D) The ability of mitochondria in diabetic organoids to reach maximal respiration is perturbed, n=8.

Autophagy is a homeostatic process that helps clear damaged organelles and mitigate cellular stress. Studies in diabetes have shown that defective autophagy plays a pivotal role in the development of diabetic complications^111,112^. However, we do not observe drastic gene differences regulating autophagy (Figure S6G). Since our data suggests mitochondrial stress is the primary driver of the defective oxidative phosphorylation, we analyzed the oxygen consumption rate using the Seahorse Flux analyzer to measure the extent of the mitochondrial stress (Figure 4D). As expected, based on the transcriptomic changes, we observed that diabetic conditions exhibited reduced basal respiration, common in diabetic individuals^113,114^. However, the most striking change was the inability of cells in diabetic conditions to reach maximal respiration when chemically stimulated with a protonophore, confirming that the mitochondria in the organoids are compromised.

### Metformin partially rescues diabetic phenotypes in diabetic salivary gland organoids

Metformin (Met) is a well-known drug given to diabetic patients to help improve insulin sensitivity by modulating AMPK signaling^115^. Though this was its original indication, it has been successfully studied in many other applications, including polycystic ovarian syndrome, obesity, prediabetes, and cardiovascular events^116^. Studies have found that metformin can improve mitochondrial function and regulate hormone levels independently of insulin^117^. Recently, studies have suggested that metformin can help mitigate salivary gland dysfunction: studies using metformin to treat patients with Sjogren’s Disease showed that treatment with metformin helps to improve disrupted calcium signaling^118^, which helped enhance salivary secretion, and others found reduced incidence of Sjögren’s Disease among metformin-taking diabetes patients compared to those who were not taking metformin^119^, suggesting that metformin can be investigated for utility in salivary hypofunction caused by inflammatory environments. We wanted to assess whether metformin could mitigate some of the diabetic phenotypes observed in SG organoids. To do this, we treated SG organoids which had been in diabetic conditions for two weeks with 50µg/mL Metformin-HCl for 24 hours (Figure 5A) and assessed the effect on diabetic phenotypes. We observed that treatment with metformin was able to partially recover the cells’ ability to reach maximal respiration (Figure 5B), as well as reduce spherical morphology and increase branching of the mitochondria (Figure 5C), suggesting that metformin treatment impacts mitochondrial fitness. Additionally, we observed that Metformin-treated samples had only one-third the number of FOXO1+ nuclei compared to untreated samples (Figure 5D), though this was still significantly higher than the osmotic controls. Moreover, we observed that Metformin treatment, at least at the dosage in this study, did not mitigate the pathogenic expression of COLIV (Figure 5E). However, it did relocalize the ductal KRT19 to the cell membrane, suggesting that treatment is helping to restore some of the organization lost in the diabetic condition. Finally, though disrupted calcium signaling in response to cholinergic agonism has been reported in Sjogren’s Disease^120^, it has not been previously evaluated in diabetic conditions. We observed that compared to control samples, diabetic conditions exhibit reduced calcium signaling in response to carbachol stimulation. However, 24hrs of treatment with Metformin partially restored this (Figure 5F).

**Figure 5:**
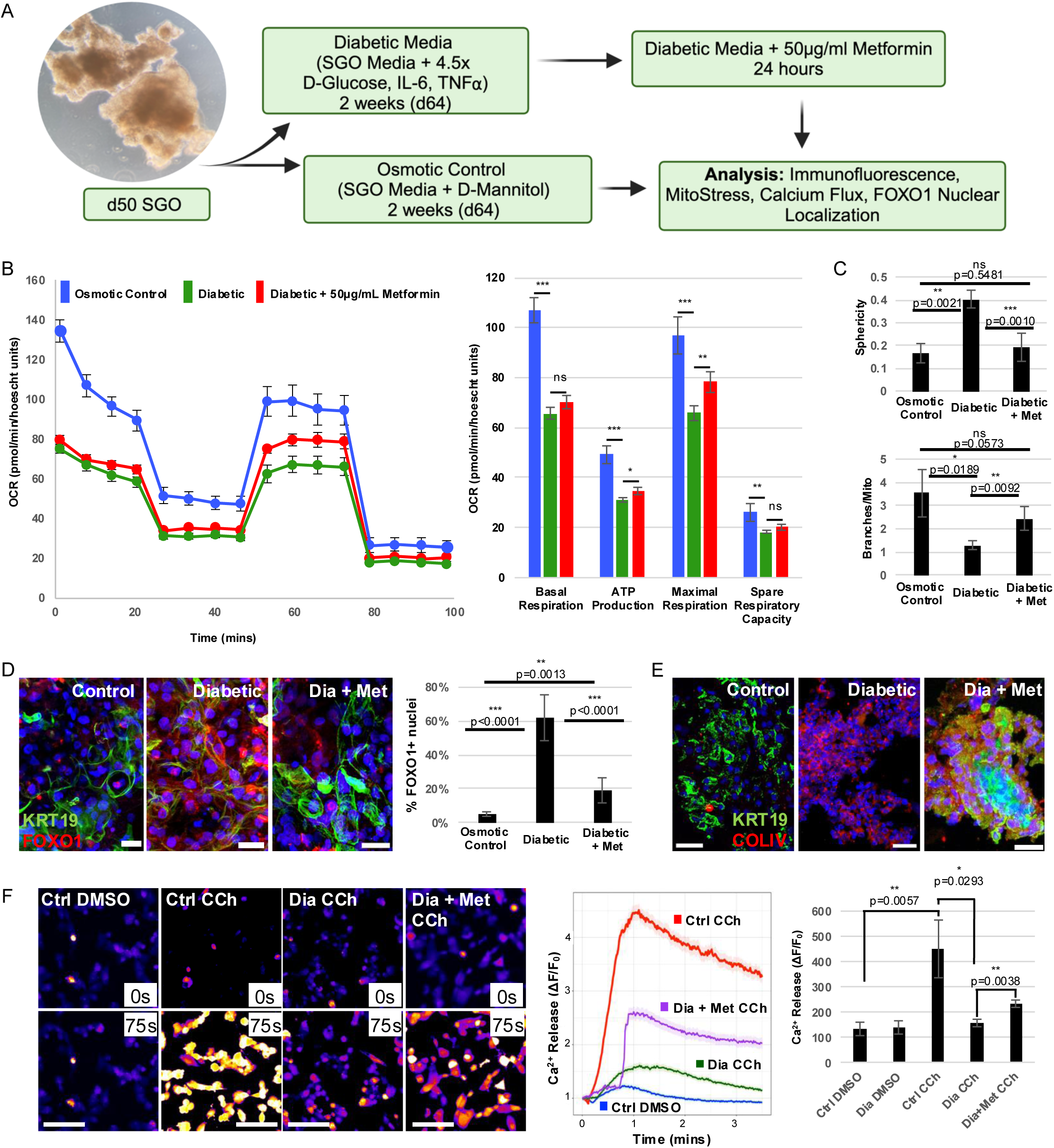
Metformin partially rescues diabetic phenotypes. (A) Schematic of treatment of organoids with metformin. (B) Metformin partially restores the ability of diabetic organoids to reach maximal respiration and (C) reduces sphericity and increases branches per mitochondria, suggesting it mitigates detrimental mitochondrial fission. (D) Metformin also partially reduces the number of FOXO1+ nuclei in the diabetic organoids. Scale bars = 25µm (E) While it does not rescue the pathogenic COLIV expression in diabetic organoids, it does restore KRT19 expression in the organoids (Scale bars = 25µm) and (F) partially rescues the calcium response to carbachol treatment.

Together, our data suggest that, in line with diabetic responses in other tissues, our salivary gland organoid model can recapitulate the consequences of a prolonged diabetic environment. These findings demonstrate that we have successfully developed a salivary gland organoid that recapitulates the basic functions of human salivary glands and proves that iPSC-derived saliva organoids can be adapted to represent salivary glands in the diabetic milieu and used to screen pharmaceuticals for therapeutic use.

## Discussion

Despite the universal acknowledgment of the need to develop permanent solutions for the various causes of salivary gland dysfunction, none have come to market. So far, there has not existed a scalable, accessible in vitro model with which to tackle questions related to numerous disease drivers or to develop effective therapeutics. While salispheres have been useful to study some aspects of regeneration and development, they have significant limitations for high-throughput studies. They rely on the derivation of human salivary gland tissue in order to procure cells. This tissue is commonly only procured in the context of salivary gland tumor excision, where surgeons take margins into healthy tissue in order to excise all the malignant tissue. Since salivary gland cancers are comparatively rare^121^, this means that these tissues are in limited supply, and the average age of patients receiving this treatment is between 50-60 years old^122^, meaning that the tissue procured may already be exhibiting hallmarks of aged salivary glands^16,17^. While existing efforts to create iPSC-derived salivary gland organoids have recapitulated salivary gland function and identity, they require extremely long culture times and technically complex microsurgery skills to create^39,123^. The long culture times and difficult techniques required for these protocols leave room for variability from person to person and make them unfit for high throughput applications. In this study, we have created a rapid salivary gland organoid protocol that provides a reliable, highly adaptable, and reproducible method by which to model genetic anomalies, explore disease etiology, screen pharmaceuticals, and develop therapeutic strategies to prevent or ameliorate current drivers of salivary gland hypofunction. As demonstrated by our work here, it is immediately useful in disease modeling applications: we adapted the organoid into a viable diabetic model that recapitulates diabetic phenotypes. Importantly, both the organoid protocol and the diabetic disease model are easily accessible to scientists of varying skill levels, making it ideal for rapid adoption in any research environment and ideal for high-throughput screening applications. Finally, with further development, this organoid has translational regenerative potential to mitigate problems like radiation-induced damage in cancer patients.

Using organoids to model diseases has improved preclinical studies leading to clinical trials in several other tissues^124–127^ and salivary gland research will be no different. Most studies seeking to understand salivary gland disease have been performed in mice due to significant morphological overlap between the species^128–132^. However, in addition to salivary gland-specific differences in innervation, structure, and time of development^133^, genomic differences in mice make it increasingly challenging to create translational solutions using mouse models^134^. Accordingly, successful clinical trials (those that have reached stage 4) have been rare and have yielded temporary solutions like oral lubricants, saliva substitutes, or secretagogues, rather than reparative therapies^135^. This organoid model provides a rapid, reliable, high-throughput tool that will immensely expand the potential for preclinical drug discovery, leading to better, more effective therapeutics in the future.

Our data showed that organoids in diabetic conditions exhibited increased expression of genes regulating focal adhesions. In normal salivary gland development, focal adhesions have been associated with promoting branching morphogenesis^141,142^, but it has never been studied in diabetic conditions. In pancreas, inhibition of focal adhesion kinase has been shown to regulate the reprogramming of pancreatic acinar cells into beta cells^143^, and focal adhesion remodeling is critical for glucose-stimulated insulin secretion^79,144^. Finally, focal adhesion overexpression has been shown to induce unintended cell migration^145^. In salivary glands, misregulating any of the pathways controlled by focal adhesions may contribute to glandular dysfunction. If signaling associated with focal adhesion kinase were involved in the transition of salivary gland progenitors to acinar cells, misregulation could perturb regeneration of damaged acinar cells. Since focal adhesion remodeling is critical for insulin secretion in the pancreas, it could also be necessary for secretory function in salivary glands. Failure to remodel could disrupt secretion, and disruption of the polarity of focal adhesions in diabetic conditions could contribute to the structural breakdown we observed in the organoids. Investigating focal adhesions’ role in diabetic salivary gland pathogenesis presents a rich new avenue of study that can be pursued with our novel diabetic salivary gland organoid model.

Notably, both oral health care and diabetes are diseases that are plagued by health disparities. Epidemiological studies demonstrate that diabetes predominantly affects certain racial and ethnic minorities and individuals with lower socioeconomic status^146–148^. They also demonstrate that oral hygiene^149,150^ and access to dental care^151–153^ are reduced for individuals of lower socioeconomic status in countries worldwide. Moreover, the disconnect between oral health care and medical care has meant that getting a diagnosis with oral health ramifications has not led to increased oral care^154,155^. This can largely be attributed to policies that do not prioritize dental care in the same way as medical care, creating policies that harm minoritized populations^151,156^. Organoid-based research can allow us to address inequities in two ways. First, iPSC-based organoids can be made from cells from diverse genetic backgrounds. This enables researchers to address racially and ethnically associated genetic differences that underpin disease susceptibility and severity, and pharmacological response. This is especially important in diseases that predominantly affect individuals of specific genetic backgrounds and can allow for the development of more effective and targeted therapeutics. Secondly, organoid-based disease models like our salivary gland model, can allow for research in tissues that cannot otherwise be studied without damaging native tissue including much more in-depth biochemical studies that can further unravel developmental and pathogenic questions regarding salivary glands.

### Limitations of Study

In this study, we created a rapid, highly adaptable, and robust salivary gland organoid model. However, our system is tempered by some broad limitations that plague organoid models including limited maturation and atypical cell organization^67^. In the current iteration, we can identify distinct duct and acinar compartments, but it will be important to conduct future studies to improve spatial organization and maturation in this model, especially as we move toward translational applications. Additionally, the current iteration of this model is unable to account for interactions of the exocrine type tissues with the immune environment. A major disease model of interest in Sjögren’s Disease, which is an autoimmune disease. The ability to create Sjögren’s Disease model or to pursue the growing interest of the role of salivary glands in viral pathogenesis will require the integration of the salivary gland organoid with the immune interface.

In vitro diabetic models like ours can provide critical data on cell behavior in diabetic conditions. However, there are some distinct differences between in vivo and in vitro diabetic models. Most basic cell culture medias have high levels of glucose. Basic DMEM/F12 has 17mM glucose, about 3 times physiological glucose. Therefore, it is common practice to use ultra-high levels of glucose, which some may argue is less clinically relevant. Therefore, as we move toward translational and clinical applications, it may be useful to reevaluate media composition and shift toward a more physiological model. However, for in vitro disease modeling and drug screening applications, our data demonstrates that the current iteration is sufficient and accurate. Additionally, though the currently used diabetic conditions do induce numerous diabetic phenotypes, it is difficult to replicate all the complexity of diabetes in vitro, including difficulties resulting from vasculopathy and neuropathy in the tissues, since our organoids lack those cell populations.

## Acknowledgments

We thank the rest of the Ruohola-Baker lab members for their helpful discussions and support throughout the study. We would like to thank Dr. Mary O’Neill and Single Cell Team at the Brotman Baty Institute for contributing to the sci-RNA-Sequencing in this work, Dr. Mary Regier of the ISCRM Genomics Core for assistance with bioinformatic analysis, and Dr. Dale Hailey and the Garvey Imaging Core for support with imaging and image processing. This work was supported by a Department of Oral Health Sciences Ruth L. Kirschstein Interdisciplinary Research Training (T90 DE021984) Postdoctoral Fellowship for D.D.E, ISCRM Fellows Program for A.P. and grants from the National Institutes of Health DE033016 (J.M., H.R.-B), DK140839 (V.C., J.M. and H.R-B), 1P01GM081619, R01GM097372, R01GM083867, NHLBI Progenitor Cell Biology Consortium (U01HL099997; UO1HL099993) SCGE COF220919 (H.R-B), and AHA 19IPLOI34760143, Brotman Baty Institute (BBI), DOD PR203328 W81XWH-21-1-0006 and Stem Cell Gift Funds for H.R-B. The Birth Defects Research Laboratory was supported by NIH award number 5R24HD000836 from the Eunice Kennedy Shriver National Institute of Child Health and Human Development.

## Author Contributions

D.D.E., J.M., and H. R-B. conceived the project and experiments. D.D.E., A.M. and A.P. performed bioinformatic analysis on sci-RNA-Seq and bulk RNA-Seq. D.D.E., A.M., and HY. J. optimized the organoid protocol and performed immunofluorescence and imaging. D.D.E. and A.P. performed the calcium response assays. D.D.E., YC. L. and Z.F. performed the Seahorse MitoStress Assays. YC. L. performed the western blots and Coomassie stains. V.C. performed EM analysis of the diabetic organoids. J.M and H. R-B. supervised the study. D.D.E., J.M., and H. R-B. wrote the manuscript with input from all authors.

## Materials and Methods

### Cell Culture

WTC-11 human induced pluripotent stem cells (Coriell, Cat# GM25256) and were maintained in 10cm culture plates coated with growth factor-reduced Matrigel (Corning, #356231) and cultured in mTeSR1 stem cell medium (StemCell Technologies, #85850) and fed daily. Cells were passaged when they reached 70-80% confluence.

### Organoid Development

To generate salivary gland organoids, passage-ready WTC-11 iPSCs were dissociated with Accutase (Stem Cell Technologies # 07920), counted, and replated in Costar Ultra Low Attachment 6 well plates (Corning, #3471) at a density of 4 × 10^5^ cells/well in mTeSR media supplemented with ROCKi (Selleck Chemical # S1049) (d-3). At each 24hrs until d0, wells were supplemented with an additional 1ml of mTeSR. At d0, spheroids are fed with Epicult-C base media (Epicult-C basal media, Epicult supplement, 1X Glutamax, 1x NEAA, 0.48ug/ml Hydrocortisone, Stem Cell Technologies # 05630) supplemented with 400nM SAG (Selleck Chemical # S7779). On d3 and d5, cells are Epicult-C base media supplemented with 400nM SAG and 50ng/mL BMP4 (R&D Systems, #314-BP-010). On d8, cells are fed with Epicult-C base supplemented with 400nM SAG, 5uM CHIR99021 (Selleck Chemical #4423), 500pM recombinant EGF (R&D Systems #236-EG), 1uM LDN-193189 (Tocris #6053), and 3.5uM NT4 (R&D Systems #268-N4). On d10, cells are fed again as on d8, but with 250ng/mL recombinant FGF10 (Peprotech #100-26). On d12, cells are switched to DMEM/F12 (ThermoFisher #11320033) containing 1x N2 supplement (ThermoFisher #17502048), 1x Glutamax (ThermoFisher #35050061), 1x NEAA (ThermoFisher #11140050), 250ng/mL recombinant FGF10, 125ng/mL recombinant FGF1 (Peprotech #AF-100-17A), and 62.5ng/mL recombinant FGF7 (Peprotech #100-19). On d15, cells are fed with DMEM/F12 containing 1x N2 supplement, 1x Glutamax, 1x NEAA, 125ng/mL recombinant FGF10, 62.5ng/mL recombinant FGF1, and 31.25ng/mL recombinant FGF7, and 300ng/mL NRG1 (Peprotech #AF-100-03). On d18 and d21, cells are embedded in ultra-low viscosity matrix of 3% Matrigel and 1% laminin (Santa Cruz Biotechnology #sc-29012) and fed with DMEM/F12 containing 1x N2 supplement, 1x Glutamax, 1x NEAA, 125ng/mL recombinant FGF10, 62.5ng/mL recombinant FGF1, 31.25ng/mL recombinant FGF7, and 500ng/mL NRG1. On d24 and d27, cells are fed with DMEM/F12 containing 1x N2 supplement, 1x Glutamax, 1x NEAA, 75ng/mL recombinant FGF10, 37.5ng/mL recombinant FGF1, 18.75ng/mL recombinant FGF7, 250ng/mL NRG1, 200ng/mL recombinant EGF, 1uM CHIR99021m 10uM SB-431542 (MedChemExpress #HY-10431), and 10nM ROCKi. From d30 to d50, cells are fed every 3 days with DMEM/F12 containing 1x N2 supplement, 1x Glutamax, 1x NEAA, 75ng/mL recombinant FGF10, 25ng/mL recombinant FGF1, 18.75ng/mL recombinant FGF7, 250ng/mL NRG1, 200ng/mL recombinant EGF, 5uM CHIR99021, and 10uM SB-431542. At every feed, plates are tilted up and organoids are allowed to sink to the bottom of the well before removing the media.

### Nuclei Extraction

Mature organoids (d56) were washed in RNase-free PBS, allowed to sink to the bottom of the tube to form a pellet. PBS was removed and the pellet was snap frozen in liquid nitrogen and stored at −80°C until enough organoids were collected to perform sequencing. Nuclei were then isolated as previously described^157^.

### Sci-RNA Sequencing

Single-cell combinatorial-indexing RNA-sequencing (sci-RNA-seq) was conducted as described previously^158^. Briefly, nuclei were thawed, permeabilized, sonicated, and distributed across 96-well plates, and tagged with the first molecular identifier using in situ reverse transcription to incorporate UMIs. Nuclei were then redistributed to other 96 well plates and tagged with a second molecular index by hairpin ligation, followed by second strand synthesis, tagmentation, purification and indexed PCR. Library was purified and sequenced on an Illumina NovaSeq platform, and aligned to GRCh38-primary-assembly, gencode.v27.

### Unbiased Clustering and Pseutotime Analysis of sci-RNA-Seq

Low quality reads were excluded from each sample by gating out cells with a UMI less than 200 and all cells with a high proportion of mitochondrial UMIs. To prevent obfuscation of cell identity in actively cycling cells, we used a Seurat vignette to regress out common cell cycle-associated genes from consideration for clustering^159^. Cells were clustered using the Monocle3 workflow^158,160,161^. Briefly, data was normalized by size factor, preprocessed using principal component analysis, dimensions were reduced using the UMAP algorithm, and clustered using unsupervised graph-based clustering analysis via the Leiden algorithm. Clusters were identified using literature-derived marker genes for salivary gland/saliva and glandular epithelium and functional characterization of highly expressed genes in each cluster.

Pseudotime analysis was performed with the Monocle3 default workflow^158^. Briefly, a machine learning technique known as reversed graph embedding is used to “learn” the principal graph and branchpoints that represent the predicted developmental trajectory and embed it back into the graph that represents the single cell dataset. Cells are assigned a pseudotime value based on their location along the predicted trajectory in relation to the root node and plotted in UMAP with coloring indicative of where a given cell falls in the biological process.

### Label Transfer from Fetal dataset

To integrate the salivary gland organoid dataset with our previously published and annotated fetal dataset, we utilized the scPoli^162^ model from the scArches2 framework^163^. The parent dataset identifier was incorporated as a batch covariate, while previously computed cell-cycle scores were included as continuous covariates for regression. Cell type annotations from the fetal dataset were provided as input labels for the scPoli prototype loss to guide the transfer learning process. For model optimization, we employed a zero-inflated negative binomial (ZINB) loss function for reconstruction within a latent space of five dimensions. The model was trained for a total of 50 epochs, including 40 pretraining epochs to establish a robust latent representation before fine-tuning on the query dataset.

### Bulk RNA-Sequencing and Analysis

Organoids were lysed and RNA was isolated using the Aurum™ Total RNA Mini Kit (Bio-Rad #7326820). RNA was quantified via A260 the NanoDrop™ One Microvolume UV-Vis Spectrophotometer, and 500ng of RNA per condition was sequenced by BGI Genomics.

Differential gene expression analysis was performed using the DESeq2 R package^164^ (version 1.46.0) in RStudio (2024.9.0; R version 4.4.0). Biotype information for genes was taken from Ensembl using the biomaRt R package (version 2.62.0) in RStudio and was used to filter the count matrix to only include protein coding genes when generating the PCA and volcano plots. The PCA plot was generated using the plotPCA function within DESeq2. The volcano plot was generated using the EnhancedVolcano R package (version 1.24.0) in RStudio and genes that passed thresholds of padj < 0.05 and log2FC > |1.5| were considered significantly differentially expressed. Heatmaps were generated using the Pheatmap R package (version 1.0.12) in RStudio. Lists of significantly differentially expressed genes were generated with DESeq2 for further analysis. KEGG analysis was performed using a graphical gene-set enrichment tool called ShinyGO^165^ and gene ontology enrichment analysis was performed using the PANTHER database^166^.

### Immunofluorescent Staining and Confocal Imaging

Salivary gland organoids intended for immunofluorescent staining were collected in an Eppendorf tube in 500µl of their warm culture media. An equal volume of 8% paraformaldehyde was added to the tube for a final concentration of 4% and fixed for a period of 15 minutes at room temperature on a nutator, then washed 3 times for 5 minutes in 1x PBS. Once fixed, organoids were stored in fresh 1x PBS at 4°C or pelleted and embedded in in O.C.T. compound (Tissue-Tek, #4583) and stored at −80°C for cryosectioning.

For those organoids to be sectioned, a Leica CM1850 Cryostat was used to create 10µm which were mounted on SuperFrost Plus slides (Fisher Scientific #12-550-15). Prior to staining, slides were washed in 1x PBS for 5 minutes to remove O.C.T compound, then overlaid in blocking solution containing 5% bovine serum albumin, 1% normal goat serum, and 0.1% Triton-X and left to incubate at room temperature for 90 minutes. After blocking, slides were overlaid with primary antibody diluted in blocking solution and placed in a humidity box in 4°C overnight. The next day, slides were washed three times in 1x PBS for 5 minutes and overlaid with AlexaFluor-conjugated secondary antibodies (Life Technologies, 1:200) or preconjugated primary antibodies diluted in blocking solution according to manufacturer recommendation and incubated at room temperature for 75 minutes. Slides were then washed four times for 5 minutes and overlaid with 5µg/mL DAPI diluted in 1x PBS and incubated at room temperature for 20 minutes. Slides were washed a final time in 1x PBS for 15 minutes, then mounted with Vectashield Hardset Antifade Mounting Medium (Vector Laboratories #H-1400) and allowed to set overnight in the dark at 4°C. After staining, slides were stored at 4°C in the dark. Slides were imaged on an inverted Nikon Eclipse Ti inverted microscope equipped with an A1R point scanning confocal system with alkali photomultiplier tubes for blue and far-red detection, and GaAsP photomultiplier tubes for green and red detection. Images were taken at 40x magnification and 1024×1024 resolution and processed using NIS Elements Advanced Research imaging software (Version 5.11.01) and Fiji ImageJ (Version 2.16.0/1.54p).

For organoids intended for whole-mount staining, organoids were placed in a 500µl centrifuge tube and submerged in Blocking Buffer containing 2.5% goat serum, 2.5% donkey serum, and 0.3% TritonX-100 in 1X PBS. Tubes are placed on a nutator and incubated overnight at 4°C. The next day, buffer is replaced with antibody dilution buffer (1%BSA and 0.3% TritonX-100 in 1X PBS) containing appropriate dilutions of primary antibodies (Table S3) and incubated overnight on the nutator at 4°C. The next day, organoids were washed 3 times for 5 minutes and submerged in antibody dilution solution containing AlexaFluor-conjugated secondary antibodies or preconjugated primary antibodies, and 5µg/mL DAPI, and incubated overnight at 4°C on a nutator. On the final day, organoids are washed 3 times for 5 minutes in 1X PBS and mounted one organoid per well of a 12 well silicone isolator (Grace Biolabs #JTR12R-A2-2.0) mounted to a glass slide (Sigma Aldrich # BI0082), in Vectashield Hardset Antifade Mounting Medium. Organoids were imaged on a Leica DMi8 confocal microscope equipped with HyD and PMT spectral detectors and LASX acquisition software [version 3.5.5IR]. Images were captured with a 40X (NA 1.1 PlanApo) water immersion lens. Images were processed with Imaris Microscopy Image Analysis Software (Version 9.9) and FIJI ImageJ Analysis software.

### Quantification of nuclear FOXO1 and AMY1A

To quantify FOXO1 nuclear localization, osmotic control, diabetic, and diabetic + metformin treated organoids were visualized by immunofluorescent staining and imaged as described above. Total nuclei were counted and nuclei that colocalized with FOXO1 stain were counted as positive. Results represented as the percent of the total nuclei that were FOXO1-positive. Results represent 5-8 samples per condition.

To quantify AMY1A, osmotic control and diabetic organoids were visualized with immunofluorescent staining and imaged as described above. Using ImageJ (Version 2.16.0/1.54p), a SUM projection of the unmodified AMY1A channel was created and the pixel intensity was measured. The diabetic and control samples were averaged and the n-fold change over the control was calculated. Results here represent the average n-fold change over control for 4 individual experiments.

### Mitochondrial Morphology Analysis

Organoids were stained with MitoTracker Orange CMTMRos (ThermoFisher #M7510) according to manufacturer protocol or with the antibody ATPB (AbCam #ab14730) and imaged as described above. To quantify changes in mitochondrial morphology due to diabetic conditions, we used a program called Mitochondria Analyzer^110^, an ImageJ plugin that uses a comprehensive pipeline for 3D quantification of mitochondrial morphology and network connectivity.

### Calcium Release Analysis

The day prior to the assay, osmotic control and diabetic salivary gland organoids were dissociated into single cells using TrypLE (ThermoFisher #12604013) according to manufacturer protocol and seeded onto Matrigel-coated 96-well (Corning, #3603) plates at 5,000 cells per well. The morning of the assay, cells were starved in a low-glucose media comprised of SILAC DMEM/F12 (A2493901), 1x N2, and 5mM D-Glucose (Sigma-Aldrich #G8270) for 6 hours. Following starvation, the cells were incubated in same low-glucose media containing 5mM Calbryte 520 AM fluorescent intracellular calcium indicator (AAT Bioquest, #20651) for 30 min at 37°C, then washed 3X with media. Just prior to image acquisition, cells were treated either with 1µM DMSO or 1µM Carbachol in DMSO. Confocal live imaging was done on a Nikon Yokogawa W1 spinning disk confocal microscope using a 20X objective air lens. Parameters for each live frame: Excitation/Emission filters for GFP fluorescence, Exposure time of 150ms, acquisition rate of 1 sec/frame, and total recording time of 5 minutes including a 30s baseline recording. Images were processed with Fiji software distribution of ImageJ 2.16.0/1.54p and frame-by-frame cellular fluorescence intensity was tracked and quantified with CellProfiler^167,168^. Condition-specific average calcium release was calculated by tracking each individual cell’s response during the recording time and computing the mean peak fluorescence achieved by all cells in the frame. An average of 50-100 cells were tracked per recording.

### Western Blot and Coomassie

For protein analysis, organoids were collected in RIPA buffer, then homogenized by repeated vortexing every 5 minutes for 20 minutes, followed by centrifugation at 14000g at 4°C for 20 minutes. The supernatant was collected. Bradford assay (Bradford reagent, Thermo Scientific #23238) was performed to obtain protein concentration (PerkinElmer Envision Multilabel Plate Reader). Standard curve was obtained using different concentrations of bovine serum albumin (BSA) (Millipore Sigma #A2153), blanked with water and PBS (average of the two). 20 ug of protein was loaded in each sample. The media in which the organoids were cultured was collected on d36 and d46 for healthy SGO, and after two weeks for osmotic control and diabetic samples, and frozen until assay was performed. Before assay, media was thawed over ice and concentrated using 30kDa molecular weight cutoff centrifuge tubes (Sigma-Aldrich #UFC9030), according to manufacturer protocol. Protein concentration was determined using Bradford Assay. Samples were mixed 1:3 with 4X Laemmli sample buffer (Bio-Rad #1610747) containing *β*-mercaptoethanol. The samples were boiled at 95°C for 10 minutes. Denatured samples were run in a 4-20% pre-cast polyacrylamide gel (Bio-Rad #4561094) in Tris-glycine-SDS running buffer at 250V for 30 minutes, then transferred onto a nitrocellulose membrane soaked in Tris-glycine-SDS with 20% methanol at 25V for 12 minutes (Trans-Blot Turbo, Bio-Rad). Membranes were blocked with 5% BSA in 1X TBS-T for 1 hour, cut and stained with primary antibodies (amylase [Cell Signaling Technology #3796] and histone-3 [Abcam #1791] or S6 ribosomal protein [Cell Signaling Technology #2217]) diluted 1:1000 in 5% BSA overnight at 4°C. Membranes were washed three times with TBST and stained with secondary antibodies (goat anti-rabbit HRP-conjugated [Bio-Rad #1706515] diluted 1:10000 for H3 and S6 membranes and 1:2000 for amylase membranes) at room temperature for 1 hour. Membranes were washed three times, then a 50-50 mixture of hydrogen peroxide and HRP substrate (Millipore #WBKLS) was added and visualization was performed on a Bio-Rad ChemiDoc Imaging system.

For Coomassie Blue staining of organoid media, 2X sample buffer (2X Laemmli buffer [Bio-Rad #1610737] 95%, *β*-mercaptoethanol 5%) was added to samples in a 1:1 ratio. The samples were boiled at 95°C for 10 minutes. Denatured samples were run in a 4-20% pre-cast polyacrylamide gel (Bio-Rad #4561094) in Tris-glycine-SDS running buffer at 250V for 30 minutes. The gel was transferred to a glass container with 200mL ultrapure water and microwaved for 60 seconds, then gently rocked at room temperature for 4 minutes. The water was discarded, and 200 mL ultrapure water was added. Two additional cycles of microwaving and rocking were performed. PageBlue protein staining solution (Thermo Scientific #24620) was added after the last wash and the gel was rocked at 20 minutes at room temperature. After staining, the gel was washed with deionized water three times. Visualization was performed on a Bio-Rad ChemiDoc Imaging system.

### Seahorse Mitochondrial Stress Test

The day prior to the assay, osmotic control and diabetic salivary gland organoids were dissociated into single cells using TrypLE (ThermoFisher #12604013) according to manufacturer protocol and seeded onto Matrigel-coated 96-well Seahorse plates at 5,000 cells per well. The Metformin-treated samples were supplemented with 50µg/mL Metformin. The morning of the experiment, the culture media was exchanged for base media (unbuffered DMEM (Sigma D5030) supplemented with 1mM sodium pyruvate (ThermoFisher #11360070) and 25-mM glucose for 1 hour prior to the assay. Substrates and selective inhibitors were injected during the measurements to achieve final concentrations of 4-(trifluoromethoxy) phenylhydrazone (FCCP, 300nM), oligomycin (5mM), and antimycin (2.5mM) + rotenone (2.5mM). The oxygen consumption (OCR) values were normalized to the number of cells present in each well, quantified by the Hoechst staining (Sigma-Aldrich #HO33342). Changes in OCR in response to substrates and inhibitors addition were defined as the maximal change after the chemical injection compared to the last OCR value before the subsequent injection.

### Ultrastructural Characterization of Salivary Gland Organoids

Salivary gland organoids were fixed in 0.1 M sodium cacodylate trihydrate buffer (pH 7.4) containing 2% (w/v) paraformaldehyde, 2.5% (w/v) glutaraldehyde, 3μM CaCl_2_ for 4 hours at room temperature, and then transferred at 4°C overnight. Samples were then post-fixed with osmium tetraoxide (1% (w/v) in H2O) and counterstained with uranyl acetate (2% (w/v) in H2O). Following gradual dehydration in ethanol, samples were embedded in Durcupan resin (Electron Microscopy Sciences, Fort Washington, PA) and polymerized overnight at 60°C as described105. Ultrathin sections (80 nm) were then cut using a 35° angle Diatome diamond knife and mounted on 300 mesh gold grids (Electron Microscopy Sciences, Fort Washington, PA). Following counterstaining with uranyl acetate (1% (w/v) in H2O) and Sato lead citrate [1% (w/v) in H2O], sections were imaged at 80 keV using an electron microscope (1200FX; JEOL, Akashima, Japan).

