## Supplemental Text and Figures for "Human iPSC-derived salivary gland organoids model diabetic salivary gland dysfunction"

We began with 2D WTC-11 iPSCs grown to 70% confluence, inducing oral epithelium by activating Hedgehog and BMP pathways through d8 and then and then repressing BMP signaling and upregulating Wnt and EGF signal for 2 days (Figure S1A). By day 10 (Figure S1B), we observed that cells had created small buds throughout the well. We then pushed salivary fate by introducing salivary epithelium driver FGF10, which we found to be expressed early in human fetal development in the mesenchyme. After two days of treatment, we observed that cell clusters exhibited more complex budded offshoots (Figure S1C) which persisted through d35. By d35, clusters had developed distinct duct-like morphology (FigureS1D) which expressed duct marker KRT19. Despite the morphological development, clusters primarily expressed duct marker KRT19, some limited KRT5 expression (Figure S1F', excretory duct marker), but no acinar markers (Figure S1E''), suggesting that FGF10 was not sufficient to induce salivary gland maturation. Because our sequencing data demonstrated that Wnt activation and TGFβ inhibition drove bifurcation of ductal progenitors toward more mature phenotypes including intercalated and striated duct, and proacinar cells (Figure S1G), we sought to modulate these pathways. To determine the appropriate concentration of both Wnt agonist CHIR99021 (Chiron) or TGFβ inhibitor SB-431542, we maintained a 1μM concentration of one while varying the other at 1μM, 5μM, or 10μM concentrations. While no concentration of either molecule was sufficient to induce the expression of acinar markers, we observed that increased concentrations of TGFβi resulted in more lumen-like structural organization (Figure S1H-J, H'-J'), while higher concentrations of Chiron disrupted cell polarization (Figure S1K-M, K'-M'). Human fetal salivary glands exhibited broad expression of *EGFR*, concentrated in excretory duct (Figure S1N, yellow circle), intercalated duct (Figure S1N, blue circle), and proacinar (Figure S1N, green circle) groups, while *EGF* was expressed primarily by striated duct (Figure S1O), suggesting that adding EGF to the protocol may help facilitate proacinar development. We observed that addition of EGF at d18 of protocol resulted in some limited expression of proacinar marker MUC4 (Figure S1Q-Q'', yellow arrows). Notably, we observed that adding EGF at d12 resulted in no MUC4 expression (not shown), suggesting that adding EGF too early is detrimental to the maturation of the cells.

Salivary glands exist in a very 3D environment. We hypothesized that growing them in a 2D environment might be perturbing their maturation. To assess how critical the transition to the 3D environment is for cell maturation, we conducted the differentiation with only FGF10, TGFβi, and Chiron (Figure S2A). We began with the 2D as described in Figure S1A and differentiated them to oral epithelium through d10. We then located the clusters that formed (Figure S1B) and used the tip of a hypodermic needle to cut these out. They were then collected and distributed 1 clump per well to an Ultra-Low attachment U-bottom 96 well plate and continued the differentiation through d35. We observed that the clumps formed spheroids and survived well, but despite limited cell death, they did not increase greatly in size. However, we found that transitioning cells to 3D (without the addition of EGF in the latter half of the maturation stage) was sufficient to induce MUC4 expression (Figure S2C') and ACTA2 expression (Figure S2C'') in addition to KRT19 (Figure S2C), suggesting that growing organoids in a 3D environment was critical to their ultimate maturation.

Studies in salivary gland development have increasingly suggested that matrix cell interactions are critical in driving morphological changes and cell fate changes. As far back as 1980, collagen synthesis was identified as critical for branching morphogenesis[1], and more recently, hydrogels linked to laminin have been show to promote salivary gland regeneration in mice[2,3] , and more recently, it was demonstrated that fibronectin dynamics in the surrounding matrix of the salivary epithelium were critical to drive clefting and branching of developing salivary buds[4], highlighting the importance of an extracellular matrix in proper salivary gland development. Many different matrices have been used in organoid environments[5], however we opted to employ Growth Factor Reduced Matrigel, which contains fibronectin, collagens, and laminins to best recapitulate the epithelial extracellular matrix. We observed some budding in the early stages of salivary gland development, so we opted to embed the organoids after the salivary gland precursor stage to capitalize on the role of the extracellular matrix protein in cell fate determination (Figure S3A). In line with other organoid models, we embedded organoids in 50% Growth Factor Reduced Matrigel. We observed early budding and branching by day 5 (Figure S3B-B') which continued to become more complex through day 11 (Figure S3C-C') and day 18 (Figure S4D-D'). However, after embedding organoids, the rigidity of the matrix obliterated all the existing buds and branches, resulting in dense, cyst-like organoids (d22, Figure S3E) which never recovered their complex structure, only sending out thin, filament like protrusions through d35 (Figure S3F-F'). This result was recapitulated in 10% Matrigel and, to a lesser extent, in 5% Matrigel (not shown), indicating that high viscosity matrices would not be amenable to salivary gland maturation. Studies in breast organoids have shown that the viscoelasticity of the extracellular environment is an important determinant of spatiotemporal tissue organization[6] and studies in gut epithelium have had success using ultra-low viscosity matrices in suspension to culture organoids[7–9], so we tried a 3% Matrigel matrix, supplemented with 1% Laminin to drive the signaling pathways we had identified through sci-Seq of human fetal tissue. Using this 3D protocol, we observed that embedding at day 18 with this matrix did not have catastrophic effects on the overall morphology of the organoids and allowed for further budding and branching that we first observe by day 5 through the remainder of the protocol (Figure 1A).

Figure S1: Modulation of Wnt, EGF, and TGFβ signals modulate structure formation and cell fate in 2D SG differentiation. (A) Schematic of experiment. (B-C) Phase images of budding from cell clumps, d10. (D) d35 2D organoids showing duct-like morphology. (E-E''') 2D organoids do not express proacinar marker MUC4 (F-F''') d35 organoids show some limited expression of basal duct marker KRT5. (G) Schematic of how Wnt and TGFβ signaling regulate cell fate in fetal salivary gland development. (H-J') Phalloidin shows that increasing TGFβ concentration increases the number of polarized clusters. (K-M') Phalloidin shows that increasing Chiron concentration decreases the number of polarized clusters. (N) Cluster map of human fetal sci-RNA-Seq showing regions of high EGFR expression. (O) Cluster map of human fetal sci-RNA-Seq showing regions of high EGF ligand expression (P) Schematic of 2D SGO protocol incorporating EGF (Q-Q'') After d35, organoids with EGF have some proacinar MUC4 expression.

Figure S2: Development of more mature SGO required 3D culture and modulation of FGF signals over time. (A) Schematic of protocol with 2D-3D transition. (B) Schematic showing the picking of d10 clusters to transition to 3D. (C'-C''') Immunofluorescence of d35 SGO showing expression of proacinar MUC4. (D) Cluster map of human fetal sci-RNA-Seq showing regions of high FGF10 ligand expression. (E) Cluster map of human fetal sci-RNA-Seq showing regions of high FGF1 ligand expression. (F) Cluster map of human fetal sci-RNA-Seq showing regions of high FGF7 ligand expression. (G) Cluster map of human fetal sci-RNA-Seq showing regions of high NRG1 ligand expression. (H) Ratio of expression of FGF10, FGF7, FGF1, and NRG1 between age groups in human fetal development.

Figure S3: 3D SG organoids exhibit structural heterogeneity and require ultra low viscosity matrix to maintain branching structures. (A) Schematic of protocol beginning in 3D with high viscosity matrix. (B-B') d5 organoidsbegin to exhibit budding and branching. (C'-C') Increased structural complexity at d11 in salivary gland organoids. (D-D') Increased budding and branching and epithelial cell polarization visible in d18 organoids prior to embedding. (E) Organoids at d22, after embedding, have lost all complex structure and exhibit cystic morphology. (F-F') Embedded organoids at d35 do not recover their complex structure and develop thing cell projections between one another. (G-I'') d50 salivary gland organoids in ultra-low viscosity matrix exhibit structural heterogeneity but always show ductal (KRT19) and acinar (AMY1A) compartmentalization.

Figure S4: Whole Western Blots and Coomassie. (A) Western Blot on salivary gland organoid protein lysate from two experiments – full blot. (B) Coomassie Stain on media collected from salivary gland organoid cultures – full blot. (C) Western blot on media collected from salivary gland organoid cultures – full blot.

Figure S5: Additional SGO Sequencing Analysis. (A) Salivary gland cluster plot and violin plot of genes commonly found secreted in saliva show that cluster 1 primarily produces these factors. (B) Plot showing expression of several pancreatic marker genes show limited to no expression, compared to known salivary gland marker (blue box).

Figure S6: Additional analysis of diabetic bulk RNA-Seq. (A) Principal component analysis of two experiments each of osmotic control and diabetic samples. (B) Heatmap of broad gene expression in the samples shows similar patterns. (C) Breakdown of genes downregulated by diabetic conditions leading to disrupted oxidative phosphorylation. (D) Heatmap showing expression pattern of differentially regulated antioxidants. (E) Heatmap showing expression pattern of differentially regulated genes associated with Mitofusion. (F) Heatmap showing expression pattern of differentially regulated genes controlling inner membrane transport. (G) Heatmap showing expression pattern of differentially regulated genes controlling autophagy.

Table S1: Antibodies used in this study

### Figure S1

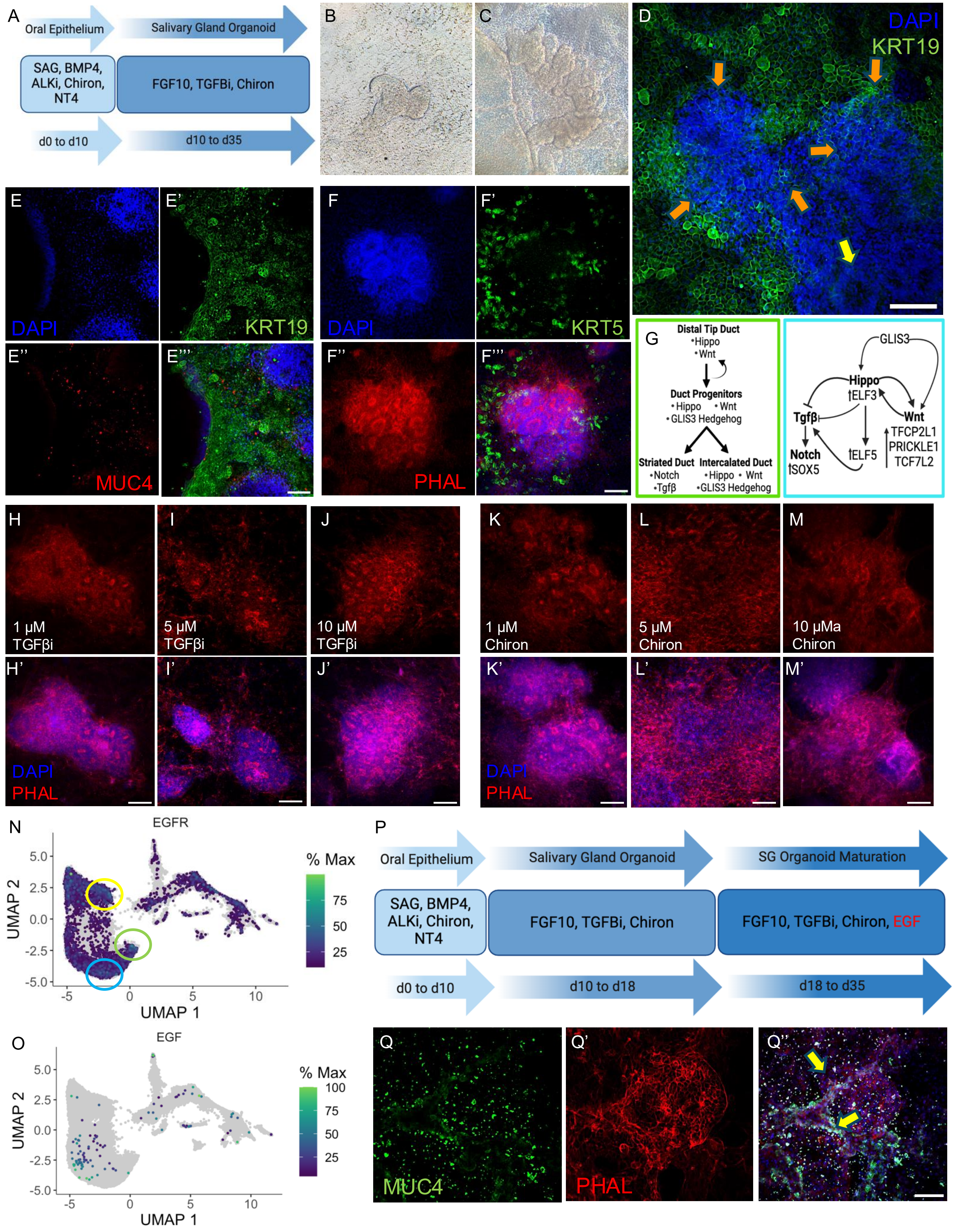
A
Oral Epithelium
Salivary Gland Organoid
SAG, BMP4, ALKi, Chiron, NT4
FGF10, TGFBi, Chiron
d0 to d10
d10 to d35
B
C
D
DAPI
KRT19
E
DAPI
E'
KRT19
F
DAPI
F'
KRT5
E''
MUC4
E'''
F''
PHAL
F'''
G
Distal Tip Duct
•Hippo
•Wnt
Duct Progenitors
•Hippo •Wnt
•GLIS3 Hedgehog
Striated Duct
•Notch
•Tgfβ
Intercalated Duct
•Hippo •Wnt
•GLIS3 Hedgehog
GLIS3
Hippo
↑ELF3
Tgfβ
Wnt
TFCP2L1
PRICKLE1
TCF7L2
Notch
↑SOX5
↑ELF5
H
1 μM TGFβi
I
5 μM TGFβi
J
10 μM TGFβi
K
1 μM Chiron
L
5 μM Chiron
M
10 μMa Chiron
H'
DAPI
PHAL
I'
J'
K'
DAPI
PHAL
L'
M'
N
EGFR
UMAP 2
UMAP 1
% Max
O
EGF
UMAP 2
UMAP 1
% Max
P
Oral Epithelium
Salivary Gland Organoid
SG Organoid Maturation
SAG, BMP4, ALKi, Chiron, NT4
FGF10, TGFBi, Chiron
FGF10, TGFBi, Chiron, EGF
d0 to d10
d10 to d18
d18 to d35
Q
MUC4
Q'
PHAL
Q''

### Figure S2

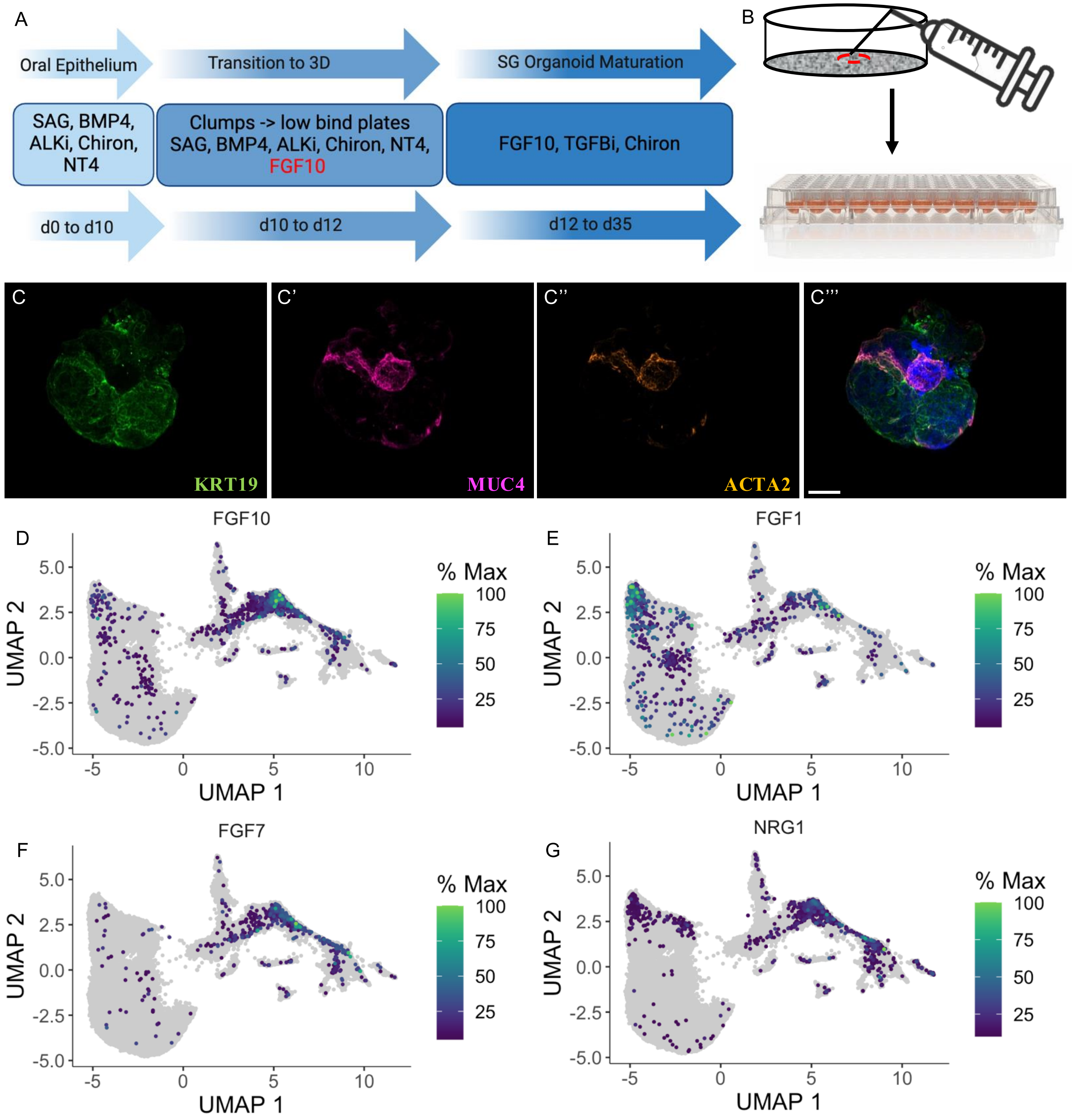


H

| | Mean Expression | | | Ratio to FGF10 | | |
|---|---|---|---|---|---|---|
| **Gene** | **12-13w** | **14-16w** | **17-19w** | **12-13w** | **14-16w** | **17-19w** |
| **FGF10** | 0.119 | 0.105 | 0.110 | 1 | 1 | 1 |
| **FGF1** | 0.059 | 0.046 | 0.037 | 0.496 | 0.438 | 0.336 |
| **FGF7** | 0.028 | 0.026 | 0.027 | 0.235 | 0.248 | 0.245 |
| **NRG1** | 0.501 | 0.555 | 0.437 | 4.21 | 5.28 | 3.97 |

### Figure S3

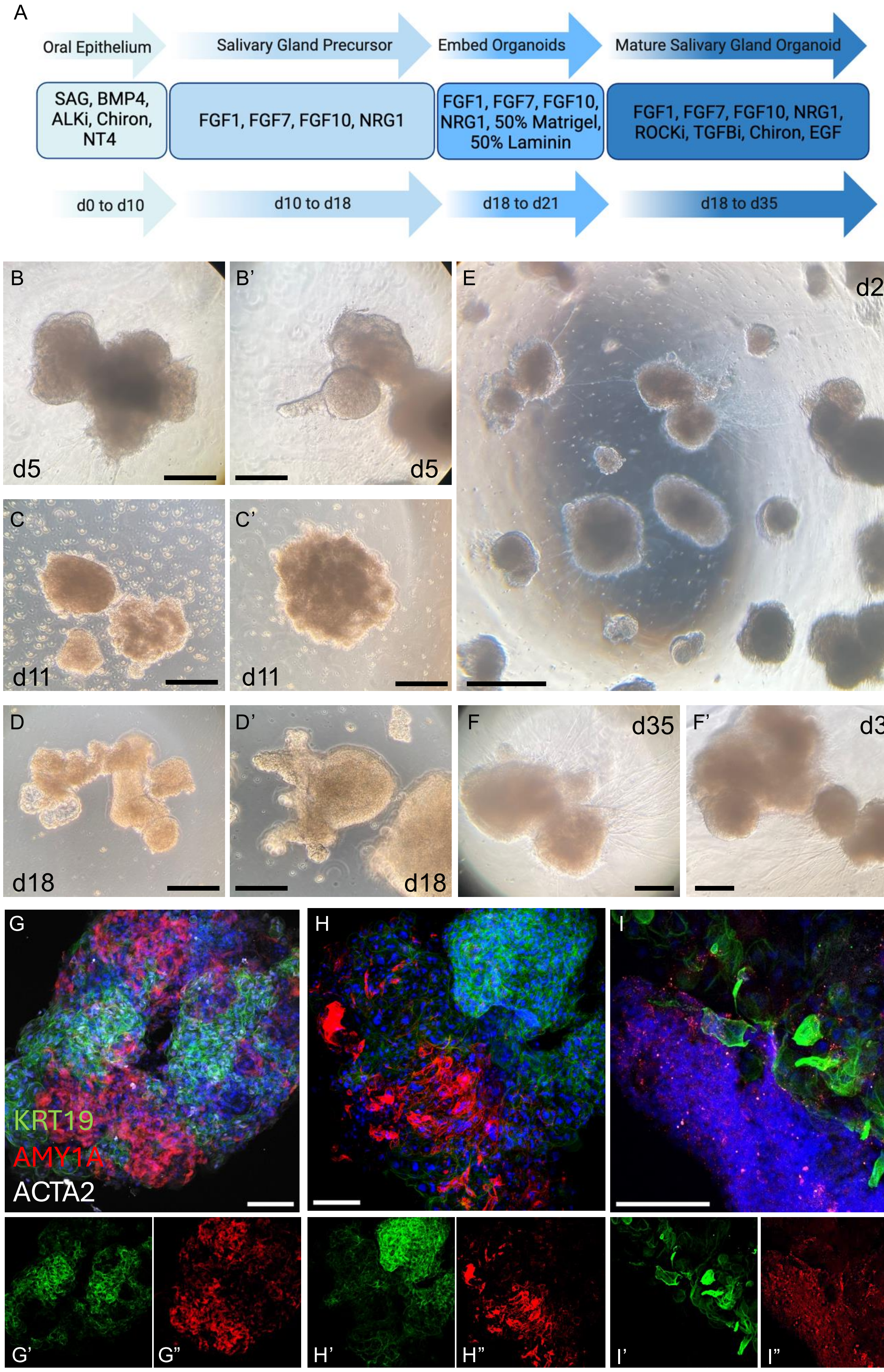
A
Oral Epithelium
Salivary Gland Precursor
Embed Organoids
Mature Salivary Gland Organoid
SAG, BMP4, ALKi, Chiron, NT4
FGF1, FGF7, FGF10, NRG1
FGF1, FGF7, FGF10, NRG1, 50% Matrigel, 50% Laminin
FGF1, FGF7, FGF10, NRG1, ROCKi, TGFBi, Chiron, EGF
d0 to d10
d10 to d18
d18 to d21
d18 to d35
B
d5
B'
d5
E
d22
C
d11
C'
d11
D
d18
D'
d18
F
d35
F'
d35
G
KRT19
AMY1A
ACTA2
H
I
G'
G"
H'
H"
I'
I"

### Figure S4

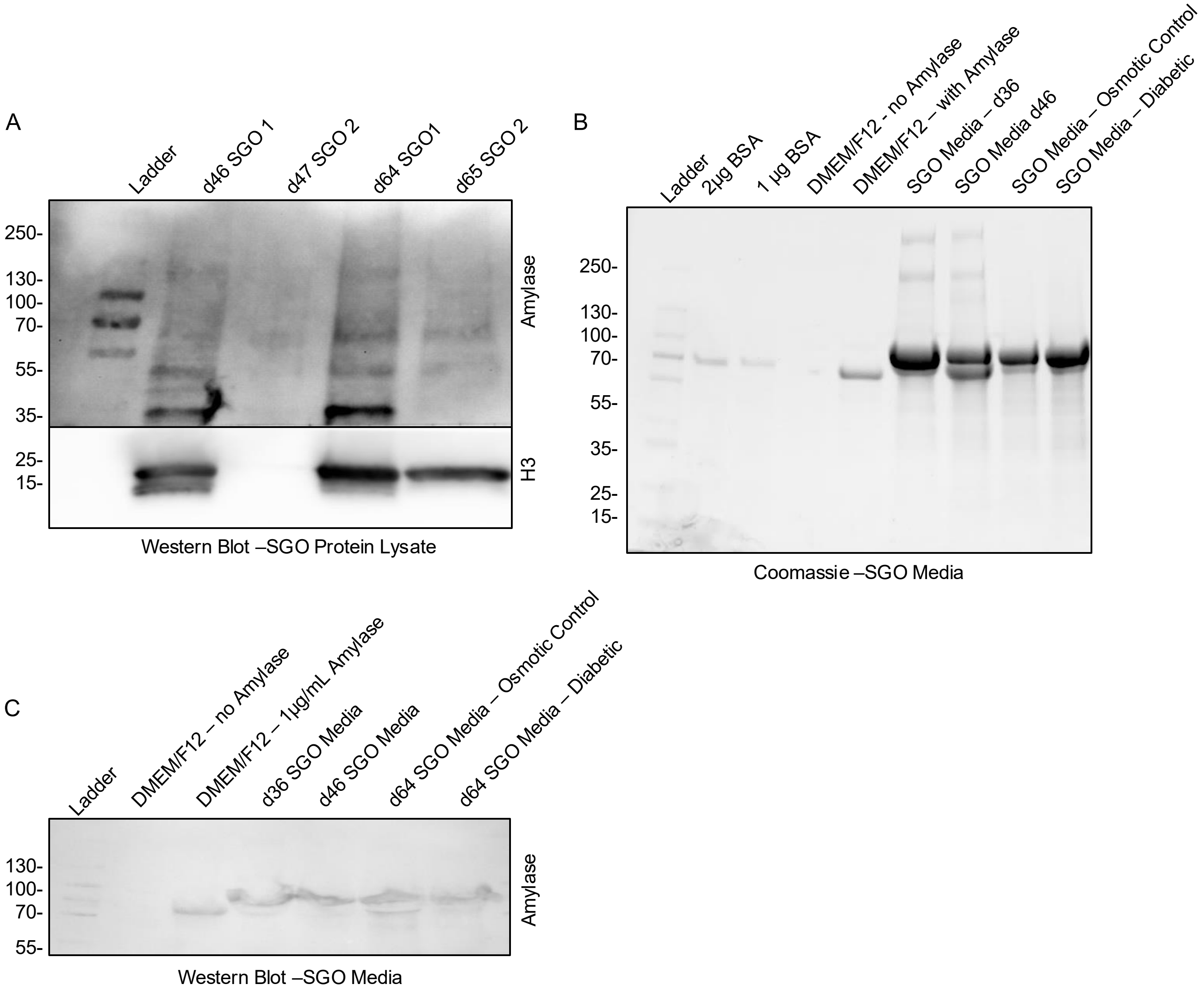

A
Ladder
d46 SGO 1
d47 SGO 2
d64 SGO1
d65 SGO 2
250-
130-
100-
70-
55-
35-
25-
15-
Amylase
H3
Western Blot –SGO Protein Lysate
B
Ladder
2µg BSA
1 µg BSA
DMEM/F12 - no Amylase
DMEM/F12 – with Amylase
SGO Media – d36
SGO Media d46
SGO Media – Osmotic Control
SGO Media – Diabetic
250-
130-
100-
70-
55-
35-
25-
15-
Coomassie –SGO Media
C
Ladder
DMEM/F12 – no Amylase
DMEM/F12 – 1µg/mL Amylase
d36 SGO Media
d46 SGO Media
d64 SGO Media – Osmotic Control
d64 SGO Media – Diabetic
130-
100-
70-
55-
Amylase
Western Blot –SGO Media

### Figure S5

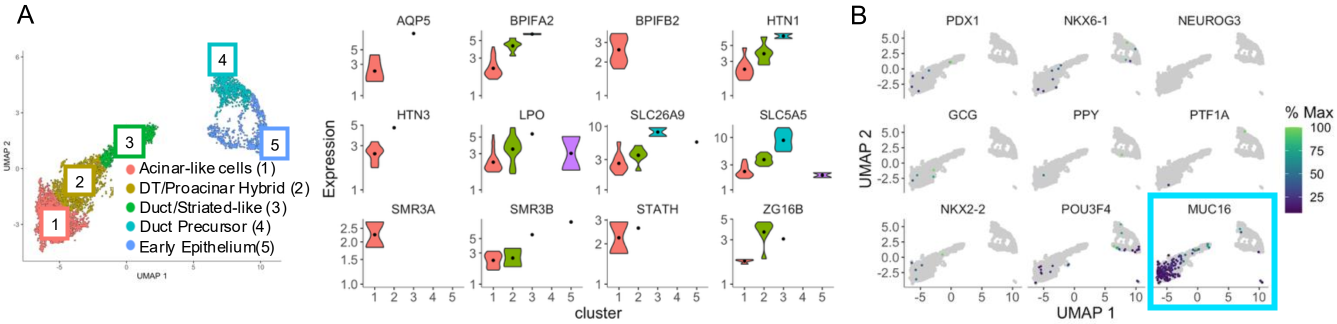

A
B
Acinar-like cells (1)
DT/Proacinar Hybrid (2)
Duct/Striated-like (3)
Duct Precursor (4)
Early Epithelium(5)
UMAP 1
UMAP 2
AQP5
BPIFA2
BPIFB2
HTN1
HTN3
LPO
SLC26A9
SLC5A5
SMR3A
SMR3B
STATH
ZG16B
Expression
cluster
PDX1
NKX6-1
NEUROG3
GCG
PPY
PTF1A
NKX2-2
POU3F4
MUC16
% Max

### Figure S6

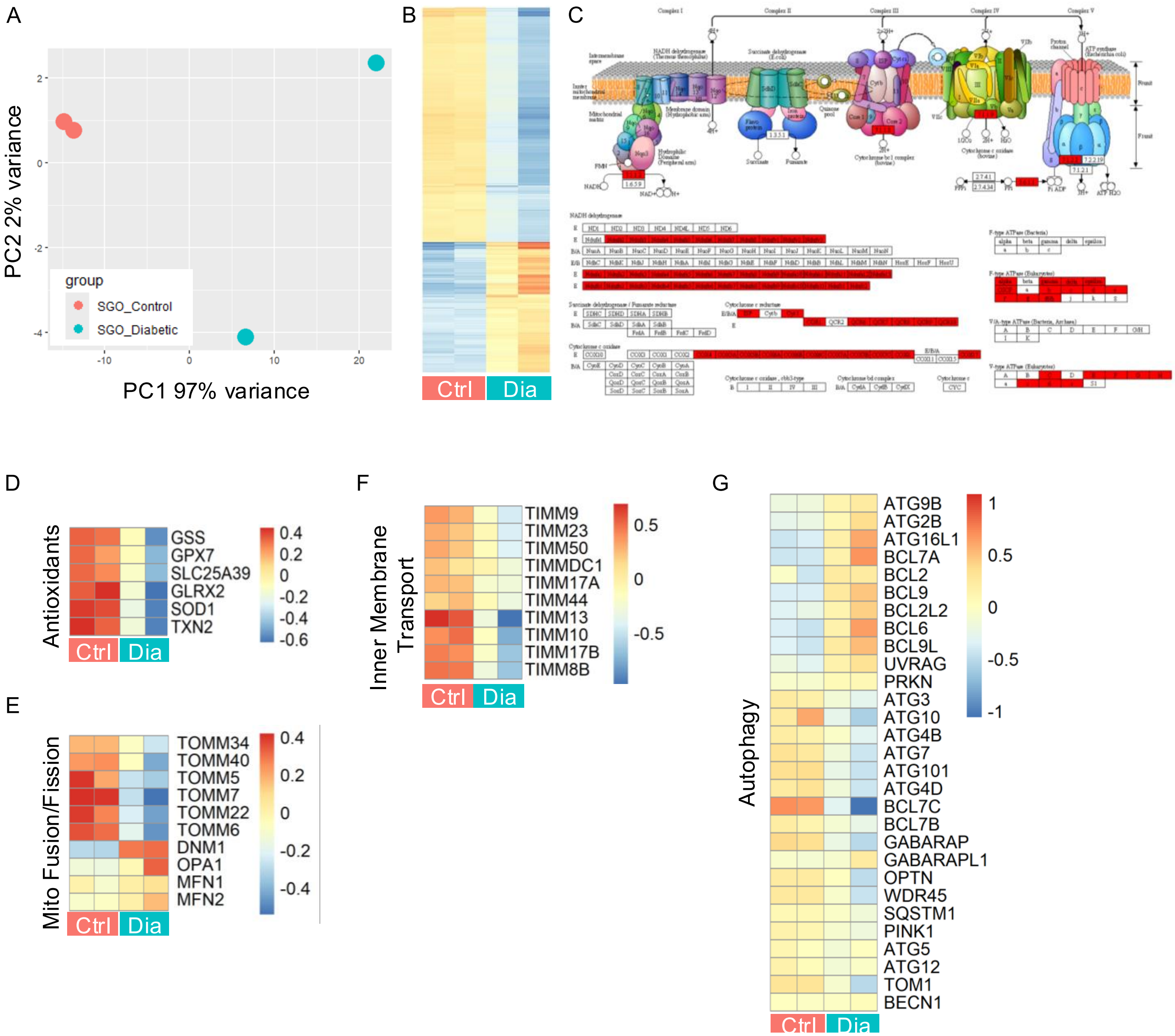
A
PC2 2% variance
PC1 97% variance
2
0
-2
-4
-10
0
10
20
group
SGO_Control
SGO_Diabetic
B
Ctrl
Dia
C
D
Antioxidants
GSS
GPX7
SLC25A39
GLRX2
SOD1
TXN2
0.4
0.2
0
-0.2
-0.4
-0.6
Ctrl
Dia
E
Mito Fusion/Fission
TOMM34
TOMM40
TOMM5
TOMM7
TOMM22
TOMM6
DNM1
OPA1
MFN1
MFN2
0.4
0.2
0
-0.2
-0.4
Ctrl
Dia
F
Inner Membrane Transport
TIMM9
TIMM23
TIMM50
TIMMDC1
TIMM17A
TIMM44
TIMM13
TIMM10
TIMM17B
TIMM8B
0.5
0
-0.5
Ctrl
Dia
G
Autophagy
ATG9B
ATG2B
ATG16L1
BCL7A
BCL2
BCL9
BCL2L2
BCL6
BCL9L
UVRAG
PRKN
ATG3
ATG10
ATG4B
ATG7
ATG101
ATG4D
BCL7C
BCL7B
GABARAP
GABARAPL1
OPTN
WDR45
SQSTM1
PINK1
ATG5
ATG12
TOM1
BECN1
1
0.5
0
-0.5
-1
Ctrl
Dia

### Table S3

| Antibody | Gene Name | Species | Dilution | Company (Catalog #) |
|---|---|---|---|---|
| KRT19 | KRT19 | Preconjugated AF488 | 1:100 (IF) | R&D Systems (IC3506G) |
| Phalloidin | N/A | Preconjugated AF 568 | 1:400 (IF) | Thermo Fisher Scientific (A12380) |
| ACTA2 | ACTA2 | Preconjugated AF 647 | 1:50 (IF) | R&D Systems (IC1420R-100UG) |
| AMY1A | AMY1A | Rabbit | 1:100 (IF)<br>1:500 (W) | Cell Signaling Technologies (3796S) |
| AQP5 | AQP5 | Mouse | 1:50 (IF) | Santa Cruz Biotechnologies (sc-514022) |
| ATPB | ATPB | Mouse | 1:100 (IF) | Abcam (ab14730) |
| NKCC1 | SLC12A2 | Rabbit | 1:100 (IF) | Cell Signaling Technologies (8351S) |
| ZO-1 | ZO-1 | Rabbit | 1:100 (IF) | Invitrogen (40-2200) |
| FOXO1 | FOXO1 | Rabbit | 1:100 (IF) | Cell Signaling Technologies (2880S) |
| MUC4 | MUC4 | Mouse | 1:100 (IF) | Thermo Fisher Scientific (35-4900) |
| KRT14 | KRT14 | Mouse | 1:100 (IF) | AbCam (ab7800) |
| Histone H3 | - | Rabbit | 1:1000 (W) | AbCam (ab1791) |
